# A single-cell atlas inclusive of age-related menopause reveals global tissue and ovarian remodeling

**DOI:** 10.1101/2025.06.20.660779

**Authors:** Hannah VanBenschoten, Zhanel Nugmanova, Audrey Olmsted, Ruixu Huang, Laasya Devi Annepureddy, Ian Gingerich, Caroline E. Kratka, Francesca E. Duncan, Olha Kholod, Brittany A. Goods

**Affiliations:** Thayer School of Engineering, Dartmouth College, Hanover NH 03755; Program in Quantitative Biomedical Sciences, Dartmouth College, Hanover NH 03755; Department of Obstetrics and Gynecology, Feinberg School of Medicine, Northwestern University, Chicago, Illinois, USA; School of BioSciences, The University of Melbourne, Parkville 3052, Australia

**Author notes:** equal contribution (Olha Kholod, Brittany A. Goods). Corresponding authors Lead contact: Brittany A. Goods.

## Abstract

Multi-tissue single-cell atlas efforts have transformed our understanding of cellular diversity across the human body and led to the creation of harmonized resources to advance research and therapeutic development. Ovarian biology, however, often remains underrepresented in these resources and is rarely analyzed with menopausal status as a biological variable. Here, we present the Menopause Cell Map (MenoMap), an integrated single-cell resource comprising more than 2 million cells from 13 healthy human tissues, including the ovary, from pre- and post-menopausal age donors. This harmonized atlas leverages curated samples from healthy female donors and enables transcriptomic comparisons across cell types, tissues, and organs while preserving menopausal status based on age as an interpretable variable. Using this resource, we show that menopause-associated gene expression changes are highly context dependent, with prominent remodeling in ovarian stromal, endothelial, immune, and reproductive cell populations. Within the ovary, post- menopausal remodeling rewired intercellular communication and shifted reproductive and steroidogenic programs toward collagen–integrin signaling, endothelial-to-mesenchymal transition, and senescence concentrated in the endothelial and stromal compartments. Cross-species comparison with young and aged mouse ovarian single-cell data showed these endothelial and stromal changes were conserved with age, most prominently in tumor necrosis factor–nuclear factor-κB signaling. Finally, we apply Human Protein Atlas-inspired specificity rules and fertility phenotype annotations to evaluate how menopausal age status affects tissue- and cell-type-specific gene classification and to prioritize ovary-enriched genes for downstream biological and translational investigation. Through this analysis, we find that the tissue specificity classification of several genes involved in reproductive-specific programs change in the pre- to post-menopausal transition, highlighting the need for age-aware and healthy donors in multi-tissue single-cell atlas efforts. Together, the MenoMap provides a human single-cell framework that leverages existing and standardized single-cell datasets curated for studying ovarian biology across reproductive aging, evaluating the influence of menopausal status on gene expression and tissue specificity, and nominating candidate genes for future investigation in reproductive biology, fertility, and target discovery.

## INTRODUCTION

The ovary is a dynamic endocrine and reproductive organ that undergoes profound cellular, molecular, and functional remodeling across the lifespan. In pre-menopausal individuals, the ovary supports follicle development, oocyte maturation, ovulation, and cyclic steroid hormone production.^1,2^ In peri- and post-menopause, follicular depletion and loss of cyclic ovarian activity are accompanied by extensive changes in endocrine signaling, stromal composition, immune state, vascular function, and tissue architecture, changes that are characterized by atrophic changes, increased inflammation, cessation of follicular activity, and reduced hormone production and responsiveness, all of which impact gene expression patterns.^3–5^ These changes have implications not only for reproductive aging, but also for systemic physiology, disease risk, and the interpretation of human tissue-specific gene expression datasets.^6–8^ Despite the central role of the ovary in reproductive and endocrine health, a comprehensive single-cell, multi-tissue atlas that explicitly incorporates both the ovary and menopausal status as an interpretable biological feature for studying female biology and driving reproductive drug discovery remains needed. More generally, reproductive transitions have critical impacts on biology and yet they are often not integrated into high-level omics analyses which removes critical context for data interpretation.

Understanding ovarian gene expression in the context of reference tissues with traceable donor age is critical for therapeutic investigation in ovarian drug development. Indeed, tissue- and cell-type-specific expression are central considerations in therapeutic target prioritization, as genes restricted to defined cellular compartments may offer greater biological specificity and lower risk of systemic effects.^9–11^ This is relevant across multiple translational contexts, including fertility preservation, ovarian aging, reproductive disease, and nonhormonal contraceptive development. However, target investigation in the ovary is complicated by the organ’s cellular heterogeneity, cyclical remodeling, and lifespan-dependent changes. ^12–15^ One of the major challenges in ovarian drug discovery occurs when genes identified to be highly expressed in the ovary or uterus are also expressed in non-reproductive tissues, or alternatively when genes are exclusive to the reproductive tract, but their disruption causes lethal or sub-fertile phenotypes.^16,17^ Genes that appear ovary-enriched in one biological context may reflect specific follicular or germ cell populations, endocrine state, or reproductive age. Conversely, genes identified in post-menopausal or bulk tissue datasets may fail to capture transcriptional programs that are specific to the functional pre-menopausal ovary.^14,18^ These issues underscore the need to better characterize ovary-specific genes in humans relative to other tissues in the body to de-risk drug discovery efforts.

Current drug discovery and gene specificity efforts rely on tissue-profiling databases such as the Genotype-Tissue Expression Project (GTEx)^19^ and Functional Annotation of Mammalian Genome (FANTOM5).^20^ While valuable, these datasets predominately include samples from post-menopausal individuals, making them unsuitable for studying reproductive function and conditions related to fertility.^21,22^ Additionally, many datasets in these databases and others rely on bulk RNA-sequencing (RNA-seq), which averages gene expression changes across entire tissues, masking cell-type-specific signals and potentially obscuring gene expression signatures of key ovarian cell populations that may be lower in abundance, such as oocytes, granulosa cells, and theca cells.^23,24^ The dynamic and heterogeneous nature of the ovary and reproductive tract necessitates a targeted data integration approach, as recent single-cell RNA-seq (scRNA-seq) studies and atlas efforts have highlighted the complex cellular composition of cycling reproductive tissues.^25–29^. scRNA-seq data can more rigorously and precisely index cell-type-specific gene expression signatures, providing rich datasets that can be used to mine specific gene signatures at various biological levels (e.g. cells, tissues, and organs). Several of these multi-tissue Atlas efforts, while extremely comprehensive with broad utility, do not include ovarian tissue,^21^ or have not focused specifically on healthy pre-menopausal and post-menopausal tissues for the use-case of drug-target identification in the ovary.^30–32^ A dedicated multi-tissue atlas that includes both pre- and post-menopausal donors as a clear variable would therefore enable direct comparison of ovarian and extra-ovarian cell types, identification of genes with reproductive-age or post-reproductive specificity, and assessment of how menopausal status alters cell-type composition, transcriptional programs, and tissue-specific gene classification.

To address this gap, we curated and integrated high-quality scRNA-seq datasets from healthy female human tissues spanning pre- and post-menopausal age donors. The resulting Menopause Cell Map (MenoMap) includes 2,010,637 single cells, 107 annotated cell types, and 13 tissues, including ovary and multiple reproductive and non-reproductive organs. By integrating these datasets in a harmonized analytical framework, we use menopausal status defined by age as an interpretable variable to examine how cellular composition, gene expression, and tissue specificity differ between pre- and post-menopausal age states. We first characterize the cellular diversity of the atlas and evaluate menopause-associated transcriptional changes across tissues and cell types. We then focus on the ovary to define menopause-associated remodeling of stromal, endothelial, immune, and reproductive cell populations, and assess whether ovarian aging signatures are conserved across human and mouse datasets. Finally, we use Human Protein Atlas-inspired specificity rules and fertility phenotype annotations to prioritize ovary-enriched genes with relevance to reproductive biology and therapeutic target investigation. Together, the MenoMap provides a human single-cell resource for studying ovarian biology across reproductive aging, evaluating the influence of menopausal status on tissue- and cell-type-specific gene expression, and nominating candidate genes for downstream biological and translational investigation. By explicitly incorporating both pre- and post-menopausal age donors, this work distinguishes features of the functional reproductive-age ovary from those associated with the post-menopausal age state, improving the interpretability of ovarian gene expression and target prioritization with healthy human datasets.

## RESULTS

### Construction and characterization of a multi-tissue single-cell atlas of pre- and post-menopausal age donors

We constructed a harmonized scRNA-seq atlas to enable systematic comparison of cellular and transcriptional programs across pre- and post-menopausal age human tissues. As much as possible we wanted to leverage existing datasets that have been published across various atlas efforts and databases. To this end, publicly available datasets were curated across 13 tissues – determined based on inclusion criteria applied to a CELLXGENE query performed in December 2025 – including ovary, fallopian tube, uterus, breast, adipose, skin, intestine, pancreas, lung, heart, liver, kidney, and eye. All metadata from source studies were maintained in the integrated scRNA-seq object, enabling retrieval of pan-atlas donor information such as age, menopause status (**Supplemental Figure 1**), health, and ethnicity (**Supplemental Data 1)**. For donors in which menopausal status was not reported, menopausal status was determined based on donor age as described in the Methods section. Interestingly, we found that skin lacked pre-menopausal samples and were highly skewed towards older donors, but the uterus was predominantly from individuals under 34 (**Supplemental Figure 1b**). Other tissues, like adipose, heart, eye, and lung were represented evenly across pre- and post-menopausal categories. Following data collection, each dataset underwent standardized preprocessing, including doublet removal, quality-control filtering based on ribosomal and mitochondrial RNA content, minimum feature detection, sample and gene filtering, removal of non-protein-coding genes, and cell-type harmonization prior to integration (**Fig. 1a, Supplemental Data 2**).^33^ The harmonized datasets were then integrated using a scVI/scANVI-based workflow, allowing donor, tissue, and dataset-level variation to be modeled within a shared latent space while preserving annotated cell identity structure for downstream analyses.^34,35^ Integration benchmarking metrics between unintegrated and final scANVI integrated layers showed preservation of biological signal with strong batch correction results across donors (**Supplemental Figure 2**).

**Figure 1.**
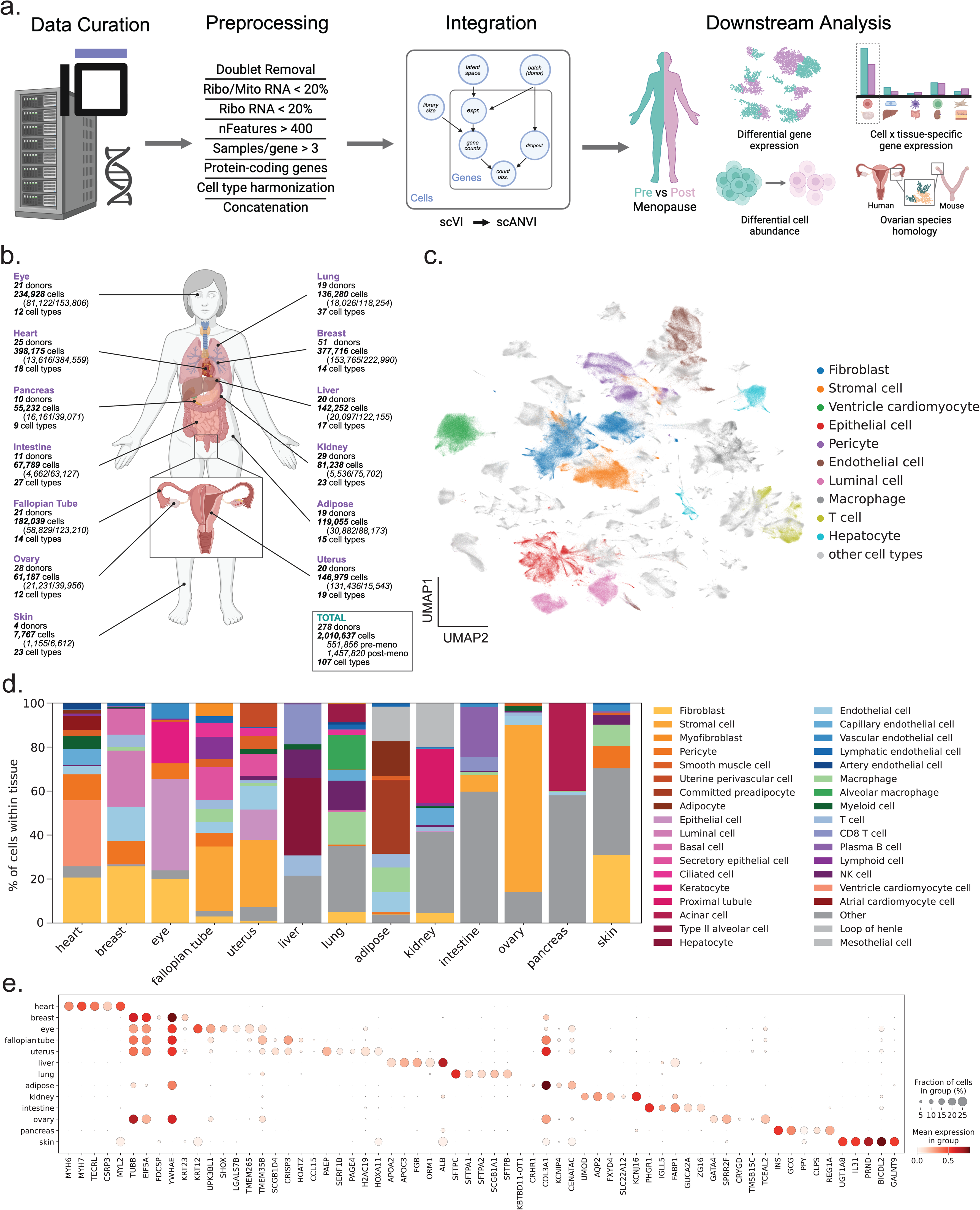
Construction and characterization of a multi-tissue single-cell atlas of pre- and post-menopausal human tissues. (a) Overview of the atlas construction workflow. Publicly available single-cell RNA-seq datasets were curated from CZ CELLxGENE across human tissues and processed through dataset-level quality control, including doublet removal, filtering based on ribosomal and mitochondrial RNA content, gene and feature thresholds, protein-coding gene retention, cell-type harmonization, and concatenation. Datasets were integrated using scVI followed by scANVI to support downstream analyses of pre- versus post-menopausal transcriptional differences, differential cell abundance, cell- and tissue-specific gene expression, and cross-species comparison of ovarian cell populations. Created in BioRender. VanBenschoten, H. (2026) https://BioRender.com/m97cjmg (b) Summary of tissues, donors, cells, menopausal status, and annotated cell types included in the integrated atlas. The final resource spans 13 tissues from 505 donors and includes over 2 million cells across 107 annotated cell types. (c) UMAP projection of the full integrated atlas colored by the top 10 most abundant cell types. Major clusters correspond to fibroblasts, stromal cells, epithelial cells, endothelial cells, immune populations, hepatocytes, cardiomyocytes, with all other tissue-specific cell types in grey, demonstrating preservation of major cellular identities after integration. (d) Stacked barplot showing the relative cell-type composition across tissues. Each bar sums to 100% of cells in that tissue and tissues are ordered by total cell number. Colors indicate cell types grouped by lineage; globally, the 35 most abundant types are shown individually and remaining types are merged into “Other.” (e) Dot plot of top tissue marker genes identified by SCANVI differential expression (each tissue versus all others, mode=’change’). For each tissue, the top five upregulated genes ranked by mean log fold change (lfc_mean) with FDR < 0.05 are shown. Rows correspond to tissues; dot color indicates mean log-normalized expression (normalize_total + log1p on raw counts), and dot size indicates the fraction of cells expressing each gene.

The resulting MenoMap is comprised of more than 2 million single cells from 278 donors, including 134 pre-menopausal and 144 post-menopausal age donors. Wherever menopausal status was not directly defined in existing study metadata, age-based inference was performed according to **Table 1** based on the approximate median age at which natural menopause occurs.^36–38^ Cell numbers varied across tissues, with particularly large contributions from breast, liver, lung, kidney, heart, and ovary datasets, while smaller but biologically relevant datasets were included for tissues such as fallopian tube, pancreas, intestine, and eye (**Fig. 1b**). This broad sampling enabled construction of a cross-tissue reference spanning both reproductive and non-reproductive organs, providing a framework to evaluate whether ovarian and menopause-age-associated transcriptional programs are tissue-restricted or shared across broader physiological systems. Visualization of the integrated atlas by UMAP demonstrated that major cell classes formed coherent transcriptional neighborhoods across the combined dataset (**Fig. 1c**). Broad populations including fibroblasts, stromal cells, epithelial cells, endothelial cells, macrophages, T cells, hepatocytes, pericytes, and additional tissue-specific populations were identifiable within the shared embedding. The presence of both shared cell types and tissue-restricted populations suggests that the integration strategy retained biologically meaningful cellular structure rather than collapsing all cells into tissue-independent clusters, as confirmed via bio-conservation metrics in integration benchmarking (**Supplemental Figure 2**). This organization supports the use of the MenoMap for both global cross-tissue comparisons across age and more focused analyses of individual cell populations.

Cell-type composition differed markedly across tissues, consistent with expected tissue architecture and specialized biological functions in each tissue (**Fig. 1d**). For example, liver contained a large hepatocyte compartment, kidney was enriched for nephron-associated epithelial populations, ovary contained a prominent fibroblast/stromal compartment, and pancreas contained a large endocrine/epithelial-associated fraction. Other tissues showed mixtures of epithelial, stromal, endothelial, immune, and smooth muscle populations. These tissue-level differences confirmed that the atlas captured expected cellular diversity across organ systems and provided a basis for comparing cell-type abundance between pre- and post-menopausal donors within each tissue. Per-tissue UMAPs and donor-wise cell composition barplots are included in **Supplemental Figure 3**. These broad compositional patterns are consistent with prior pan-tissue and cell-type-resolved human atlas efforts, which have shown that major cell classes, including epithelial, endothelial, stromal, immune, and tissue-specialized parenchymal populations, retain recognizable transcriptional identities across organs while also displaying organ-specific differences in abundance and gene expression.^21,22,39^

Cell-type marker analysis further confirmed that the integrated atlas preserved tissue-and cell-type-specific transcriptional programs (**Fig. 1e**). Dot plot visualization highlighted canonical marker genes with enriched expression in expected tissues and cellular compartments, including tissue-restricted expression patterns such as myocyte-associated *MYH6* and *MYH7* in heart, corneal-associated *KRT12* in eye, hepatocyte-associate *ALB* in liver, and alveolar *SFTPC* in lung.^40–43^ Of note, not all tissue-level marker genes have strong, known associations with tissue-specific cell types; for instance, *TUBB*, *YWHAE,* and *EIF5A* are expressed ubiquitously across several tissues, but known to have particularly high expression in brain and neurological compartments.^22^ These ambiguous marker genes could reflect confounding from gaps in reference tissues or limitations of marker gene identification at the tissue, rather than cell-type, level. Together, these results demonstrate that the integrated atlas captures broad cellular diversity across pre- and post-menopausal human tissues while retaining biologically interpretable cell identity and tissue-specific gene expression patterns.

### Menopause-associated transcriptional changes are cell-type and tissue specific across the integrated atlas

We next evaluated whether pre- and post-menopausal cells occupied distinct transcriptional states within the MenoMap. Visualization of the full atlas by menopausal status showed broad overlap between pre- and post-menopausal cells across the shared UMAP space, indicating that global cell identity was largely preserved across groups rather than being dominated by menopausal status (**Fig. 2a**). However, tissue-level visualization revealed more localized shifts in cell-state distribution (**Fig. 2a**, **Supplemental Figure 4**). In the ovary, pre-and post-menopausal age cells showed clear regional separation within several transcriptional neighborhoods, suggesting that menopause is associated with pronounced ovarian cell-state remodeling. In contrast, breast cells showed more extensive overlap between groups, consistent with more subtle or cell-type-specific transcriptional changes rather than a broad tissue-wide shift.

**Figure 2.**
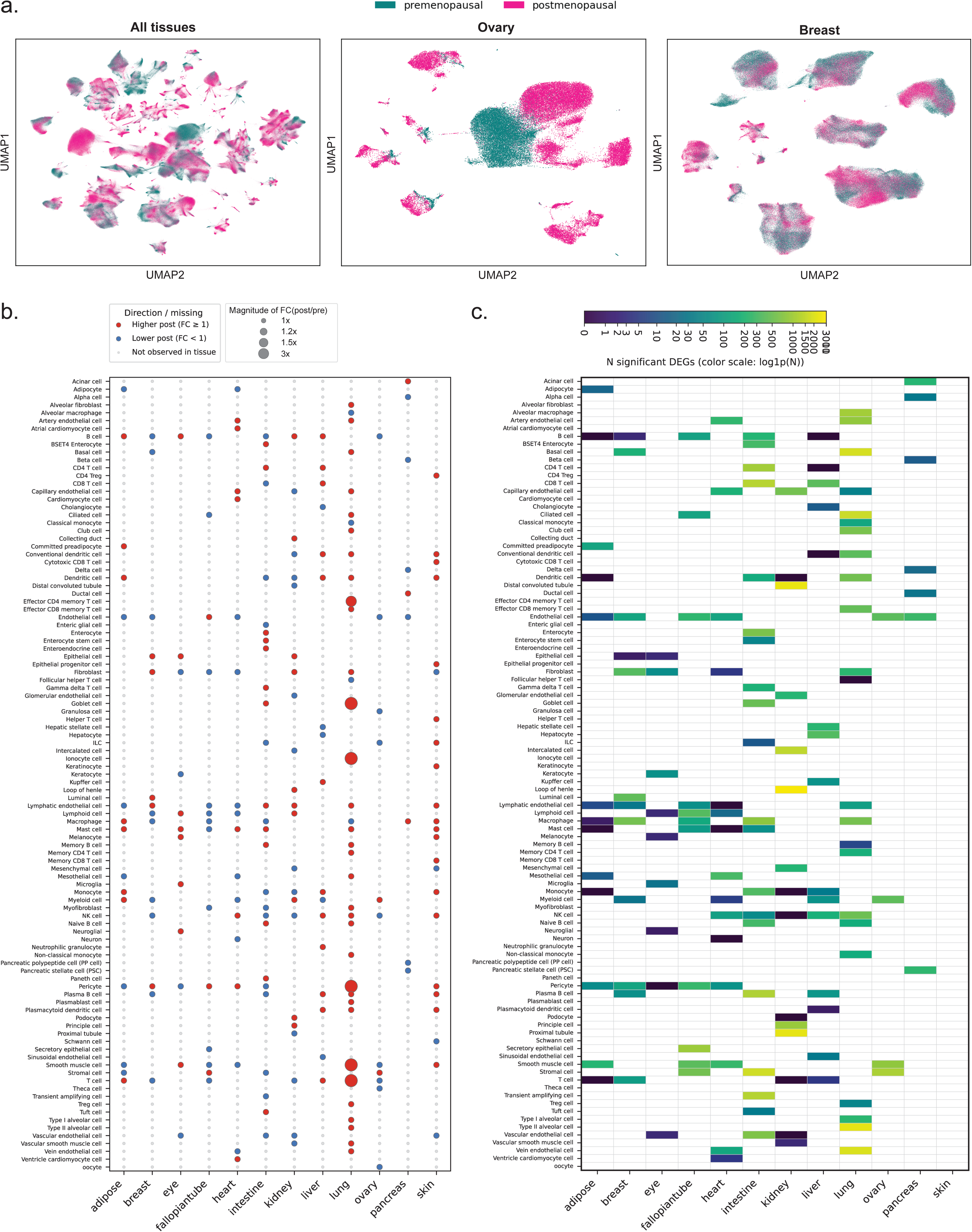
Menopause-associated transcriptional remodeling is tissue- and cell-type-specific across the integrated single-cell atlas. (a) UMAP visualization of cells from all tissues (left), ovary (middle), and breast (right) colored by menopausal status (pre-menopausal = teal; post-menopausal = pink). (b) Dotplot summarizing menopause-associated shifts in harmonized cell-type composition across tissues. Each dot indicates the postmenopausal-to-premenopausal fold change (FC) in the proportion of a given cell type within a tissue, computed from all premenopausal and postmenopausal cells pooled within that tissue. Red indicates higher postmenopausal proportion (FC ≥ 1); blue indicates lower (FC < 1); dot size reflects FC magnitude. Light grey dots mark cell types not observed in that tissue. Asterisks denote tissue-cell-type pairs with credible menopause-associated compositional effects by scCODA (inclusion-based FDR = 0.1; default). Rows, harmonized cell types; columns, tissues. (c) Heatmap showing the number of significantly differentially expressed genes (DEGs) between premenopausal and postmenopausal cells for each harmonized cell type (rows) within each tissue (columns). For each tissue-cell-type pair with at least 15 premenopausal and 15 postmenopausal cells, DEGs were identified using donor-aware pseudobulk-based differential expression with pyDESeq2. A gene was counted as significant if it passed P_adj_ < 0.05 and had an absolute mean log fold change > 1. Color intensity indicates the DEG count on a log1p scale (viridis colormap). Unshaded cells indicate pairs with insufficient cells or failed DE.

We next asked whether menopause was associated with shifts in cell-type composition across tissues. For each tissue containing both pre- and post-menopausal age cells, we calculated the proportion of each harmonized cell type within each menopause group and estimated post- versus pre-menopausal fold changes in relative abundance (**Fig. 2b**). This analysis revealed tissue- and cell-type-specific differences across tissues, with some cell populations showing increased relative abundance and others showing decreased relative abundance in post-menopausal donors. These patterns were not uniformly distributed across tissues, but instead appeared concentrated in specific tissue–cell-type combinations, suggesting that menopause-associated compositional changes are context dependent rather than global across all organs. To determine whether these abundance shifts were robust to donor-level variation and the compositional structure of single-cell data, we applied a single cell decomposition data analysis framework (scCODA) separately within each tissue using donor-level samples as replicates and performed unpaired, cross-sectional comparisons across menopausal status.^44^ Interestingly, we found that no tissue-cell-type pairs reached scCODA credibility at the target FDR threshold, indicating that the observed pooled fold-change patterns should be interpreted as descriptive trends rather than statistically supported differential abundance. This lack of significant compositional effects may reflect limited power within individual tissues, donor-to-donor heterogeneity, uneven sampling across menopause groups, or modest changes in cell-type abundance relative to transcriptional state changes. Together, these results suggest that menopause-associated differences in the MenoMap are more strongly supported at the level of cell-state and gene-expression remodeling than by robust shifts in major cell-type proportions.

To identify the cell populations contributing to transcriptomic differences between pre-and post-menopause age groups, we performed differential expression analysis within each cell type and tissue. Across tissues, significant menopause-associated differential expression was detected in stromal, epithelial, endothelial, immune, and tissue-specialized cell populations (**Fig. 2c**). The direction and magnitude of these changes varied substantially by tissue and cell type, with some populations showing higher expression in post-menopausal cells and others showing lower expression relative to pre-menopausal cells. Notably, several ovarian cell populations, including stromal and epithelial-associated compartments, displayed relatively large numbers of differentially expressed genes and larger fold-change magnitudes, which is expected in the context of menopause. Other tissues, including lung, liver, pancreas, breast, and skin, also contained cell types with significant pre- versus post-menopausal differential expression, indicating that menopause-associated transcriptional remodeling is not restricted to reproductive tissues. These differences include shifts in alveolar cells in the lung, and immune cells in the intestine. Together, these results suggest that menopause is associated with coordinated but highly context-dependent transcriptional remodeling across several cell types. The strongest effects appear in specific cell populations within select tissues, with the ovary showing particularly prominent cell-state and gene-expression differences between pre- and post-menopausal donors among ovarian endothelial, stromal, and myeloid cells.

### Menopause remodels intercellular communication networks and transcriptional programs in ovarian stromal and endothelial cells

To characterize how the menopausal transition reshapes intercellular signaling within the ovary, we analyzed premenopausal and postmenopausal ovarian tissue from the MenoMap and annotated cell types using canonical markers (Supplementary Figure 5e,f; see Methods). The two states differed markedly in composition (Supplementary Figure 5a–d): premenopausal ovaries retained many ovarian cell types, including the follicular compartment (oocyte, granulosa, theca) and diverse immune populations, albeit with substantial inter-donor variability, whereas postmenopausal ovaries have stromal-dominated populations (stromal, endothelial, smooth muscle, and myeloid), with consistent loss of the follicular and lymphoid compartments across donors. We then performed cell–cell communication analysis using CellChat on each subset and integrated the results with differential gene expression and gene set enrichment (**Fig. 3**). To distinguish intrinsic signaling remodeling from this loss of cellular diversity, we focused the differential and enrichment analyses on stromal and endothelial cells, which were present in both states.

**Figure 3.**
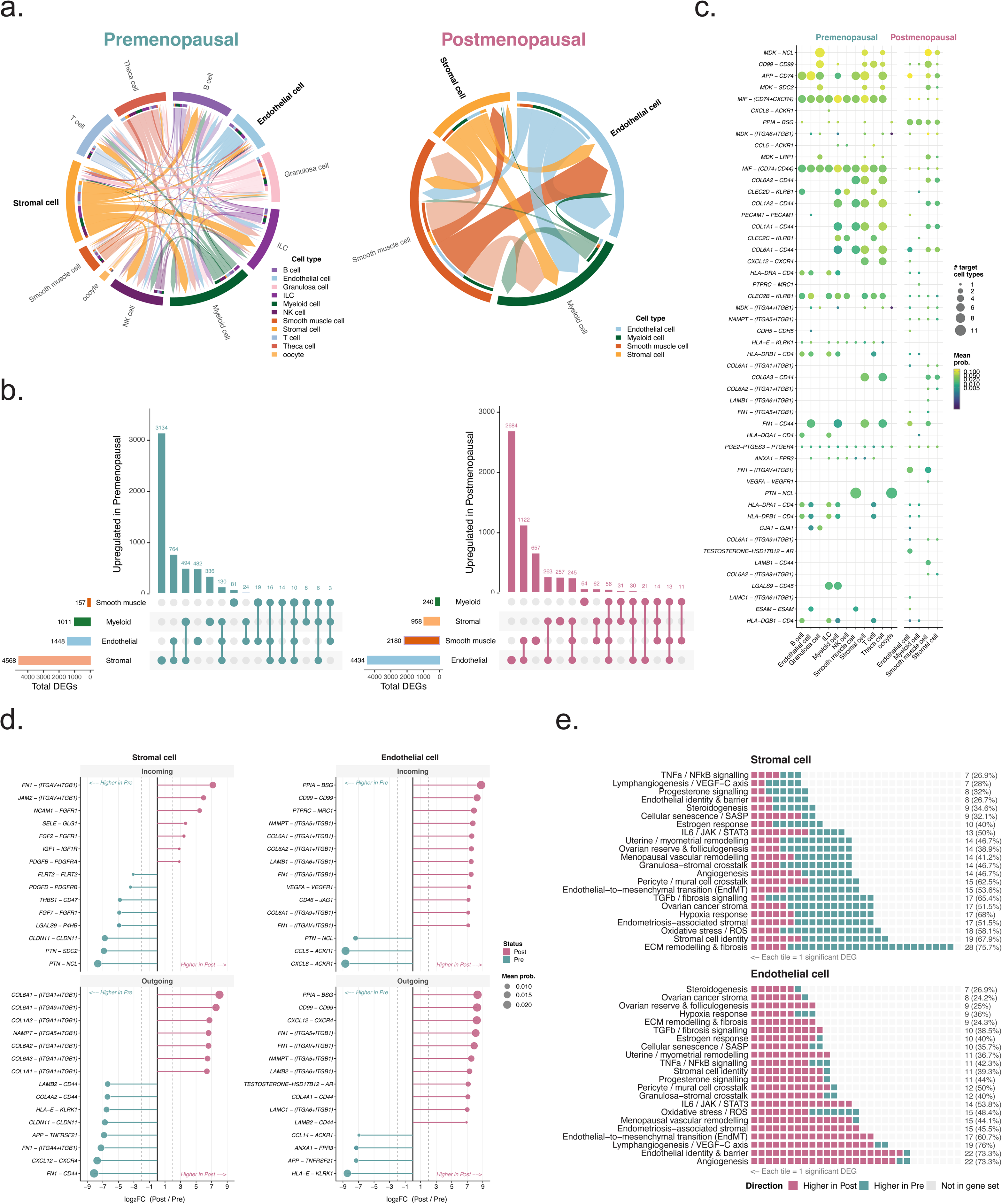
Rewiring of intercellular communication across the menopausal transition. (a) Chord diagrams depicting the global landscape of ligand–receptor-mediated cell–cell communication inferred by CellChat in premenopausal (left) and postmenopausal (right) human ovarian tissue. Arc width is proportional to the aggregate interaction probability; cell types are color-coded as indicated. (b) UpSet plots quantifying the number of differentially expressed genes (DEGs; adjusted *p* < 0.05, |log^2^FC| > 0.25) upregulated in the premenopausal (left) or postmenopausal (right) state across major ovarian cell types. Horizontal bars denote the total DEG count per cell type; vertical bars show the size of intersecting gene sets. (c) Bubble plot of the top 50 differentially active ligand–receptor (LR) pairs across all sender–receiver cell-type combinations, ranked by mean communication probability. Bubble size encodes mean probability; color indicates the number of target cell types engaged. Pairs enriched in the premenopausal or postmenopausal state are shown side by side. (d) Lollipop plots depicting differential LR interaction strengths (log_2_FC, postmenopausal vs. premenopausal) for the top incoming (upper) and outgoing (lower) interactions in stromal cells (left) and endothelial cells (right). Dot size reflects mean communication probability; blue denotes enrichment in the premenopausal state, red in the postmenopausal state. (e) Waffle-tile enrichment maps displaying the overlap between cell-type-specific DEGs and curated biological gene sets for stromal (upper) and endothelial (lower) cells. Each tile represents one significant DEG; red tiles indicate higher expression postmenopause, blue tiles higher expression premenopause. Enrichment significance was assessed by hypergeometric test with Benjamini–Hochberg correction; annotation at right denotes adjusted *p*-value (*, ** or ***) and the number and percentage of gene-set members represented.

We first characterized intercellular communication across menopausal status (**Fig. 3a**). In the premenopausal ovary, chord diagrams revealed a dense communication network including all major ovarian cell types, such as granulosa cells, theca cells, stromal cells, endothelial cells, smooth muscle cells, myeloid cells, T cells, NK cells, ILC, B cells, and oocytes. By contrast, the postmenopausal ovary exhibited fewer interactions, with communication largely restricted to stromal, endothelial, smooth muscle, and myeloid cells, reflecting a profound loss of reproductive cell types and their associated signaling networks. This transition mirrors the documented follicular depletion and immune remodeling that accompany ovarian aging.^8,15,45^ To quantify the transcriptional changes underlying this communication shift, we next performed differential gene expression analysis between premenopausal and postmenopausal ovarian cell types (**Fig. 3b**). UpSet plots indicated that stromal cells exhibited the most extensive transcriptional differences between menopausal groups. Using our predefined differential-expression criteria, 4,568 genes showed higher expression in premenopausal stromal cells, whereas 4,434 genes showed higher expression in postmenopausal stromal cells. Endothelial cells displayed the second-largest transcriptional shift, with 1,448 genes expressed at higher levels in premenopausal samples and 2,180 genes expressed at higher levels in postmenopausal samples. Because these totals include genes with a broad range of effect sizes, subsequent analyses focused on genes and pathways showing the most robust and biologically meaningful differences. Notably, the postmenopausal endothelial compartment had a higher number of upregulated genes relative to the premenopausal state, suggesting that vascular remodeling is a prominent feature of the postmenopausal ovary.^46^ Taken together, these findings demonstrate that the postmenopausal ovary undergoes extensive transcriptional remodeling, particularly within stromal and endothelial compartments.

We next examined the specific ligand–receptor pairs underlying differential intercellular communication in stromal and endothelial cells (**Fig. 3d**). Analysis of the top differentially active interactions in both incoming and outgoing directions revealed several major communication axes that were altered across menopausal status. In the premenopausal state, stromal cells exhibited increased outgoing signals through collagen–integrin signaling axes (COL1A1, COL3A1, COL5A1, COL6A2–(ITGA9+ITGB1)), PTN–NCL, and NAMPT–(ITGA5+ITGB1), among others, consistent with active extracellular matrix (ECM) deposition and growth factor signaling.^47^ Incoming premenopausal interactions into stromal cells were dominated by FN1–(ITGA5+ITGB1) and FGF–FGFR pathways, reflecting active fibroblast growth factor signaling and fibronectin-mediated cell adhesion.^48^ Conversely, the postmenopausal stromal cell exhibited increased incoming interactions from CXCL8–ACKR1, CCL5–ACKR1, and FN1–CD44, together with shifts in outgoing collagen–integrin signals, suggesting a proinflammatory and ECM-remodeling phenotype.^49^ A similar pattern was observed in endothelial cells, where postmenopausal upregulation of TESTOSTERONE–HSD17B2–AR, VEGFA–VEGFR, and MIF-related interactions was noted, pointing to altered steroidogenesis and angiogenic remodeling in the postmenopausal vasculature.^50^

To put these interaction-level changes in a broader biological context, we assessed the representation of curated gene sets within DEGs specifically in stromal and endothelial cells (**Fig. 3e**). In stromal cells, gene sets for ECM remodeling and fibrosis, stromal cell identity, and endometriosis-associated stromal were among the most represented, with a predominant higher-in-postmenopausal direction. This was accompanied by enrichment of TGFβ/fibrosis signaling, hypoxia response, and ovarian cancer stroma gene sets, suggesting a shift toward a fibrotic, stress-responsive stromal phenotype after menopause. By contrast, premenopausal enrichment was seen in gene sets related to ovarian reserve and folliculogenesis, granulosa-stromal crosstalk, progesterone signaling, and steroidogenesis – signatures consistent with the active reproductive function of the premenopausal ovary.^29,51^ In endothelial cells, the most significantly enriched postmenopausal gene sets included angiogenesis, endothelial identity and barrier, lymphangiogenesis/VEGF-C axis, and endothelial-to-mesenchymal transition (EndMT), while premenopausal enrichment was observed for steroidogenesis, ovarian cancer stroma, and ovarian reserve signatures. These findings collectively show a coordinated transcriptional and intercellular communication shift in the menopausal ovary, wherein, as expected, the reproductive and steroidogenic programs of the premenopausal state change to fibrotic, inflammatory, and vascular remodeling programs in the postmenopausal ovary. The MenoMap also highlights CCIs, such as CXCL12–CXCR4 and PTN–NCL, that may be important in the context of ovarian aging.

### Menopause alters tissue and cell-type specificity of gene expression across the MenoMap

To assess whether there are shifts in tissue- and cell-type specificity of gene expression across menopause status, we classified genes using Human Protein Atlas-inspired cell-type specificity rules (**Fig. 4a**).^22^ Genes were assigned to four categories: cell type enriched, group enriched, cell type enhanced, or low specificity, based on the relative expression of each gene across annotated cell populations. This enabled comparison of whether genes gained or lost cell-type specificity between pre- and post-menopausal age groups. Overall, the distribution of genes across specificity categories was broadly similar between pre- and post-menopausal age donors, but a subset of genes changed specificity class with menopause (**Fig. 4b**). Most genes were classified as either low specificity or cell type enhanced in both groups, while smaller numbers were classified as group enriched or cell type enriched. In the pre-menopausal age group, 5,242 genes were classified as low specificity, 11,364 as cell type enhanced, 157 as group enriched, and 154 as cell type enriched. In the post-menopausal age group, these categories contained 5,149, 11,483, 152, and 133 genes, respectively. Although the global distribution of specificity classes was largely preserved across pre and postmenopausal age donors, the movement of individual genes between categories suggests that menopause may be associated with remodeling of cell-type-restricted gene expression programs rather than global disruption of gene specificity.

**Figure 4.**
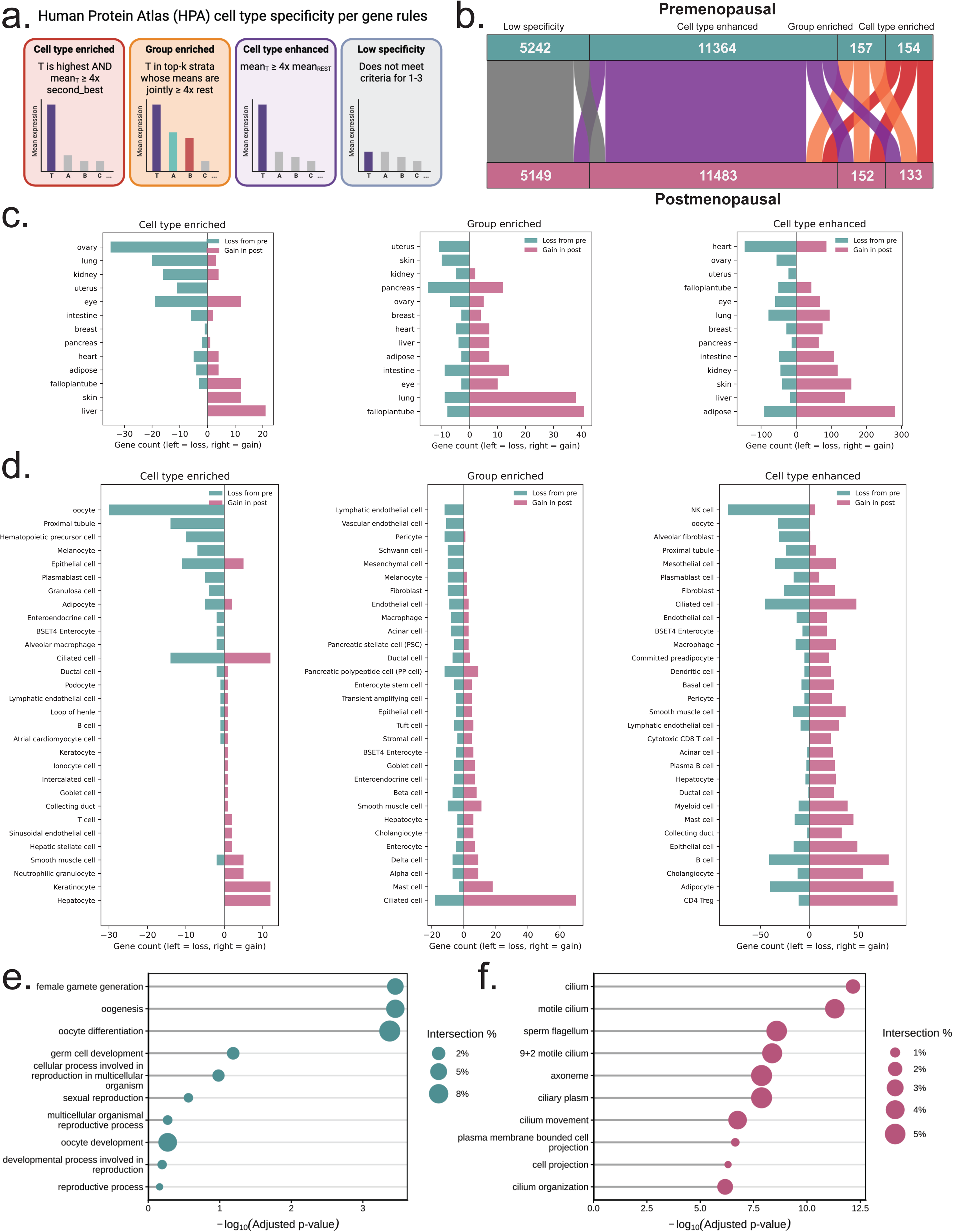
Menopause-associated remodeling of tissue and cell-type-specific gene expression programs. (a) Overview of human protein atlas-defined gene specificity rules. Genes are defined as not detected, cell type enriched (top stratum ≥4-fold above second), group enriched (top 2–10 strata jointly ≥4-fold above remainder), or cell type enhanced (≥4-fold above mean of others), or low specificity (detected but do not meet last three criteria). Created in BioRender. VanBenschoten, H. (2026) https://BioRender.com/xaq59y5 (b) Alluvial flow diagram showing pre- to post-menopausal transitions of individual genes between cell type enriched (red), group enriched (orange), cell type enhanced (purple) and low specificity (grey) criteria. (c) Tissue-level gains and losses in specificity-associated genes across the cell type enriched (left), group enriched (middle), and cell type enhanced (right) categories. Negative values (teal bars) indicate genes lost from the corresponding specificity class in the post-menopausal atlas, while positive values (pink bars) indicate genes gained in the post-menopausal atlas. (d) Cell-type-level gains and losses in specificity-associated genes across the cell type enriched (left), group enriched (middle), and cell type enhanced (right) categories. Negative values (teal bars) indicate genes lost from the corresponding specificity class in the post-menopausal atlas, while positive values (pink bars) indicate genes gained in the post-menopausal atlas. Each panel shows the top 30 cell types ranked by total gain + loss magnitude for one high-specificity category. (e) Dot plot showing the top five significantly enriched Gene Ontology (BP/MF/CC) or terms from g:Profiler over-representation analysis using the gene set from pre-menopausal ovary cell-type enriched genes. Dot size represents the percentage of genes in each term represented by the query set, and the x-axis shows −log10 adjusted p-value. (f) Dot plot showing the top five significantly enriched Gene Ontology (BP/MF/CC) or terms from g:Profiler over-representation analysis using the gene set from ppst-menopausal fallopian tube group enriched genes. Dot size represents the percentage of genes in each term represented by the query set, and the x-axis shows −log10 adjusted p-value.

We next examined which tissues contributed to gains or losses in gene specificity (**Fig. 4c**). Changes were not uniformly distributed across tissues, suggesting that menopause-associated shifts in specificity are tissue dependent. For cell type enriched genes, several reproductive and endocrine-associated tissues showed changes, including ovary, uterus, fallopian tube, and breast, along with non-reproductive tissues such as liver, lung, kidney, adipose, skin, intestine, and eye. Group enriched and cell type enhanced genes showed broader tissue-level redistribution, with some tissues gaining specificity-associated genes in the post-menopausal atlas and others losing them. These results suggest that menopause may alter the degree to which genes are restricted to particular tissues or cellular compartments, with effects extending beyond the ovary alone. This also suggests that menopausal status of donors in multi-tissue cell atlases has a meaningful effect on specificity classifications; therefore, attention must be paid to inclusion of donors across the lifespan to capture accurate specificity classification in the context of both biology and drug target evaluation.

At the cell-type level, gains and losses in specificity were concentrated in distinct cellular compartments (**Fig. 4d**). For cell type enriched genes, oocytes showed the strongest loss of specificity-associated genes in the post-menopausal age donors, consistent with the expected depletion or transcriptional remodeling of the ovarian germ cell compartment after menopause and, critically, the fact that no post-menopausal oocytes were retained in post-menopausal donors in this atlas limited direct comparisons here.^52–56^ Additional changes were observed across granulosa cells, epithelial cells, ductal cells, hepatocytes, keratinocytes, and other tissue-specialized populations. Group enriched and cell type enhanced genes showed changes across a broader range of stromal, epithelial, endothelial, immune, and tissue-specialized cell types, including ciliated cells, basal cells, endothelial cells, macrophages, smooth muscle cells, hepatocytes, and pancreatic-associated cell populations. Together, these findings indicate that menopause-associated remodeling of gene specificity occurs at both the tissue and cell-type levels and this is broadly distributed across tissues.

Gene ontology enrichment analysis provided biological context for genes with altered specificity. Genes that were cell type enriched in ovarian tissue in pre-menopause and lost post-menopause age groups were enriched for reproductive and oocyte-associated processes, including female gamete generation, oogenesis, oocyte differentiation, germ cell development, and cellular processes involved in reproduction (**Fig. 4e**). In contrast, genes that gained group enriched status in fallopian tubes in post menopause were enriched for cilia-related processes, including cilium, motile cilium, axoneme, ciliary plasm, cilium movement, and cilium organization (**Fig. 4f**). These enrichments suggest that menopause-associated shifts in cell-type specificity are linked to biologically coherent programs, including germ cell identity and ciliated epithelial functions. Together, these analyses show that while the overall landscape of gene specificity remains broadly conserved between pre- and post-menopausal age donors, menopause may be associated with remodeling of tissue- and cell-type-specific expression programs that are quite broad. The strongest signal includes loss of oocyte-associated specificity in the ovary, supporting the interpretation that menopause alters not only differential gene expression, but also the cellular restriction of gene expression across the MenoMap.

### Conservation of ovarian aging signatures across human and mouse

Given that we aimed to support the identification of candidate therapeutic targets for reproductive health across pre- and postmenopausal states, it is important to determine which human age-associated changes can be modeled in mice. Mouse models remain central to mechanistic studies and preclinical drug development, yet differences in ovarian cellular composition and gene expression may limit the direct translation of findings across species. We therefore first compared two previously published mouse ovarian single-cell datasets to evaluate their cellular composition, annotation consistency, and suitability for integration with the human atlas (**Supplementary Fig. 6**). Despite differences in the relative abundance of individual populations, both mouse datasets contained the major ovarian cell types and showed broadly consistent expression of cell-type-specific markers. Stromal cells were among the most abundant populations in both datasets, whereas endothelial cells were represented at lower but comparable frequencies, supporting the use of these two compartments for cross-dataset and cross-species comparisons.

To determine whether the transcriptional and signaling changes observed between the human pre- and postmenopausal ovary are conserved across species, we next integrated the human single-cell data with ovarian single-cell data from young and aged female mice from these prior studies (**Fig. 5**). This cross-species analysis focused on stromal and endothelial populations because they were consistently represented across the human and mouse datasets and were among the cell types showing the strongest menopause-associated transcriptional and intercellular communication changes in the human ovary.

**Figure 5.**
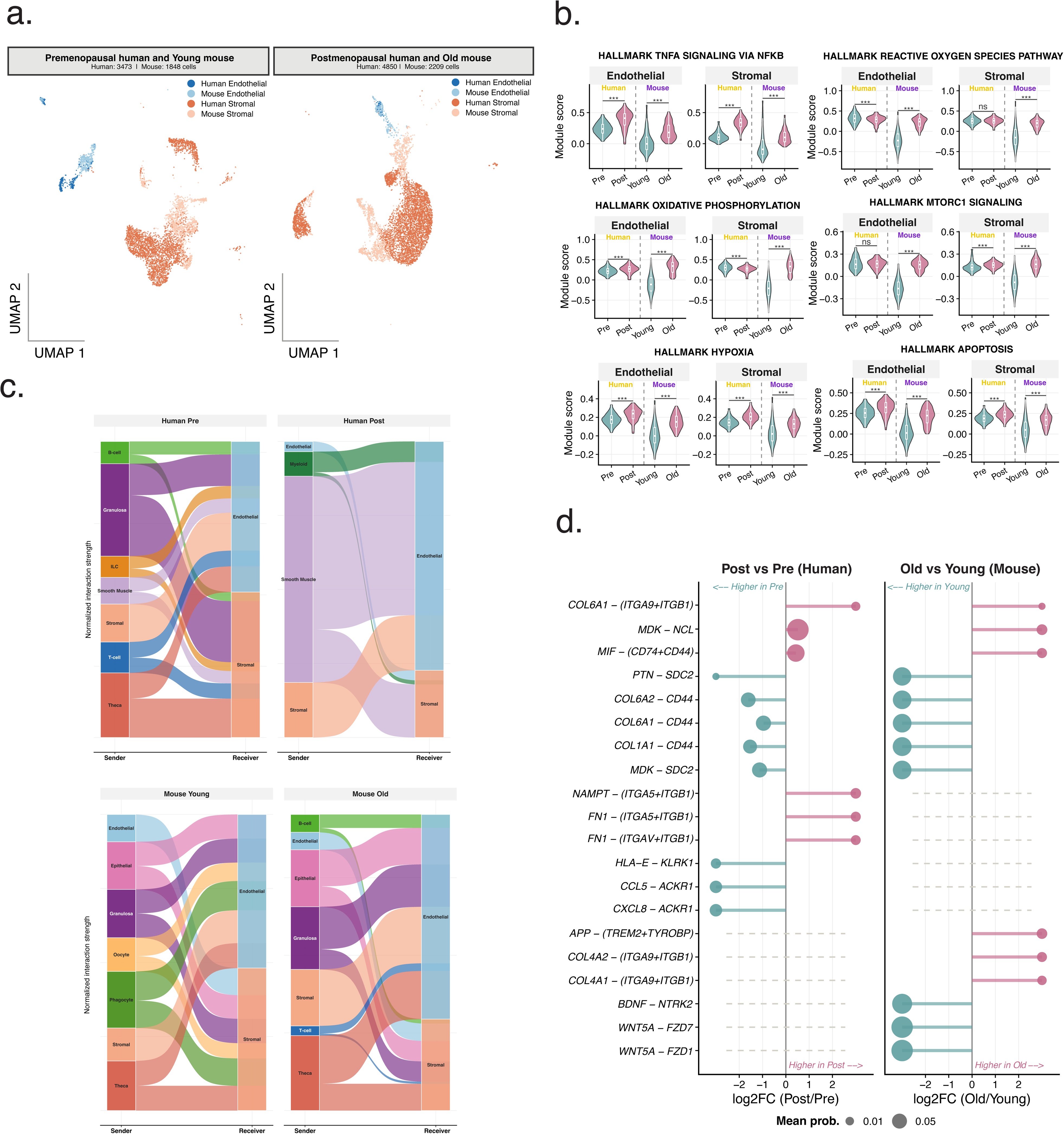
Cross-species conservation of ovarian ageing programs in stromal and endothelial compartments. (a) UMAP projections of integrated human and mouse ovarian single-cell transcriptomes, shown separately for the premenopausal human / young mouse cohort (left; *n* = 3,473 human, 1,630 mouse cells) and the postmenopausal human / old mouse cohort (right; *n* = 4,850 human, 2,209 mouse cells). Cells are colored by cell type (Endothelial, Stromal) and species (Human, Mouse). (b) Violin plots of module scores for four hallmark ageing programs: steroidogenesis, cellular senescence, oxidative stress and TGF-β/fibrosis, which were computed for endothelial (left) and stromal (right) cells in premenopausal/postmenopausal human and young/old mouse datasets. Statistical significance was assessed by Wilcoxon rank-sum test (*** *p* < 0.001; ns, not significant). (c) Alluvial (Sankey) diagrams of normalized cell-to-cell interaction strength, illustrating the rewiring of signaling flow among ovarian cell types in human premenopausal, human postmenopausal, mouse young and mouse old conditions. Ribbon width is proportional to normalized interaction strength between sender (left) and receiver (right) cell populations. (d) Lollipop plots comparing differential LR interaction strengths in human (postmenopausal vs. premenopausal, left; log^2^FC) and mouse (old vs. young, right; log_2_FC) for selected LR pairs. Pairs are stratified as conserved-gained, conserved-lost or species-specific. Dot size encodes mean communication probability. For conserved pairs, Pearson correlation coefficients between human and mouse log_2_FC values are indicated.

UMAP projections of the integrated human and mouse datasets, colored by cell type and species, demonstrated broad transcriptional similarity between ovarian stromal and endothelial cells across species (**Fig. 5a**). In the premenopausal human and young mouse comparison (human, 3,473 cells; mouse, 1,848 cells), human and mouse stromal and endothelial populations occupied partially overlapping regions of the integrated UMAP space, consistent with conservation of their core transcriptional identities. Similar integration was observed between postmenopausal human and aged mouse cells (human, 4,850 cells; mouse, 2,209 cells), although greater separation of some populations suggested the presence of species-specific features of ovarian aging.

To assess the conservation of age-associated biological programs more systematically, we calculated module scores for all 50 MSigDB Hallmark gene sets in stromal and endothelial cells. Figure 5b presents the pathways showing the most pronounced differences between reproductive-age groups. Several metabolic and stress-response programs, including reactive oxygen species, oxidative phosphorylation, mTORC1 signaling, hypoxia, and apoptosis, exhibited significant age-associated changes, although the direction and magnitude of these effects varied by species or cell type. In contrast, TNFα signaling via NF-κB was consistently elevated in both postmenopausal human and aged mouse stromal and endothelial cells. Notably, this was the only canonical inflammatory pathway among the most strongly differential Hallmark programs displayed, suggesting that activation of TNFα/NF-κB signaling may represent a particularly conserved feature of ovarian aging. Together, these results support the utility of mouse models for investigating selected components of human ovarian aging while also emphasizing that the conservation of individual transcriptional programs should be evaluated on a pathway- and cell-type-specific basis.

We next analyzed the global architecture of cell-cell communication in each condition. Interestingly, unlike our prior analyses that showed globally-altered CCI networks with age in the ovary in humans likely driven by compositional shifts in cell types, we mostly found that the total number of incoming and outgoing signals young and old mice were maintained (**Fig. 5c**). For example, we identified several signals being sent from immune cells in both old and young mice to endothelial or stromal cells. Interestingly, this included an emergent population of T cells and B cells in old mice that we did not see in postmenopausal ovaries from humans. We also found that the total number of interactions driven by smooth muscle cells in postmenopausal ovaries was large, which we did not recapitulate in old mouse ovaries. Taken together, these data highlight compositional and CCI network shifts that occur in human and mouse, and suggest that there may be some species-specific differences due either to species or to differences in aging trajectories.

To identify the specific ligand-receptor interactions most strongly altered with aging in the ovary, we constructed a comparative dot plot of the top differentially active communication pairs in human (post vs. pre) and mouse (old vs. young) stromal and endothelial cells (**Fig. 5d**). The log_2_ fold-change plots revealed a set of interactions consistently increased in the premenopausal/young condition and decreased in the postmenopausal/old condition across both species. Notable premenopausal/young-enriched interactions included COL6A1–(ITGA9+ITGB1), MDK–NCL, MIF–(CD74+CD44),^57^ and COL6A2–CD44, all of which showed concordant downregulation in both post-human and old-mouse conditions. Conversely, WNT5A–FZD7 and WNT5A–FZD1 were consistently increased in the post/old condition in both human and mouse, nominating these interactions as conserved drivers of postmenopausal ovarian remodeling. Additional interactions conserved in the postmenopausal/old direction included APP–(TREM2+TYROBP) and BDNF–NTRK2, suggesting conservation of neurotrophin and amyloid precursor protein signaling axes during ovarian aging. Together, these findings establish a cross-species framework of conserved and divergent signaling remodeling in the aging ovary, providing a foundation for mining putative drug targets.

### Prioritization of ovary-enriched reproductive genes using cell-type specificity and fertility phenotype evidence

We next wanted to use the MenoMap to identify genes that may be important in the context of pre- and post-menopause and represent a degree of druggability to better enable translational studies based on human datasets.^11,58^ To this end, we prioritized ovarian genes with potential relevance to reproductive biology and specifically for contraception or fertility. We compared gene expression in each ovarian cell type against all other ovarian and extra-ovarian cell populations restricted to pre-menopausal age donors. Genes were classified using Human Protein Atlas-inspired specificity rules as described above, including cell type enriched, group enriched, cell type enhanced, low specificity, and not detected categories (**Fig. 6a**). This workflow enabled identification of genes with ovarian cell-type-restricted expression patterns in premenopausal age donors. Several genes showed highly restricted expression in ovarian cell populations, with the strongest cell-type-specific signal observed in oocytes (**Fig. 6b**). Canonical oocyte-associated genes^55^, including *NLRP5, NLRP14, FIGLA, KPNA7, NLRP9, DPP3, ZP4, ZP5,* and *PADI6*, displayed high average expression and a large fraction of expressing cells in the oocyte compartment, with little or no expression across stromal, granulosa, epithelial, endothelial, myeloid, smooth muscle, T cell, B cell, and NK cell populations. This pattern supports the ability of the atlas-based specificity framework to recover expected ovarian germ cell markers.

**Figure 6.**
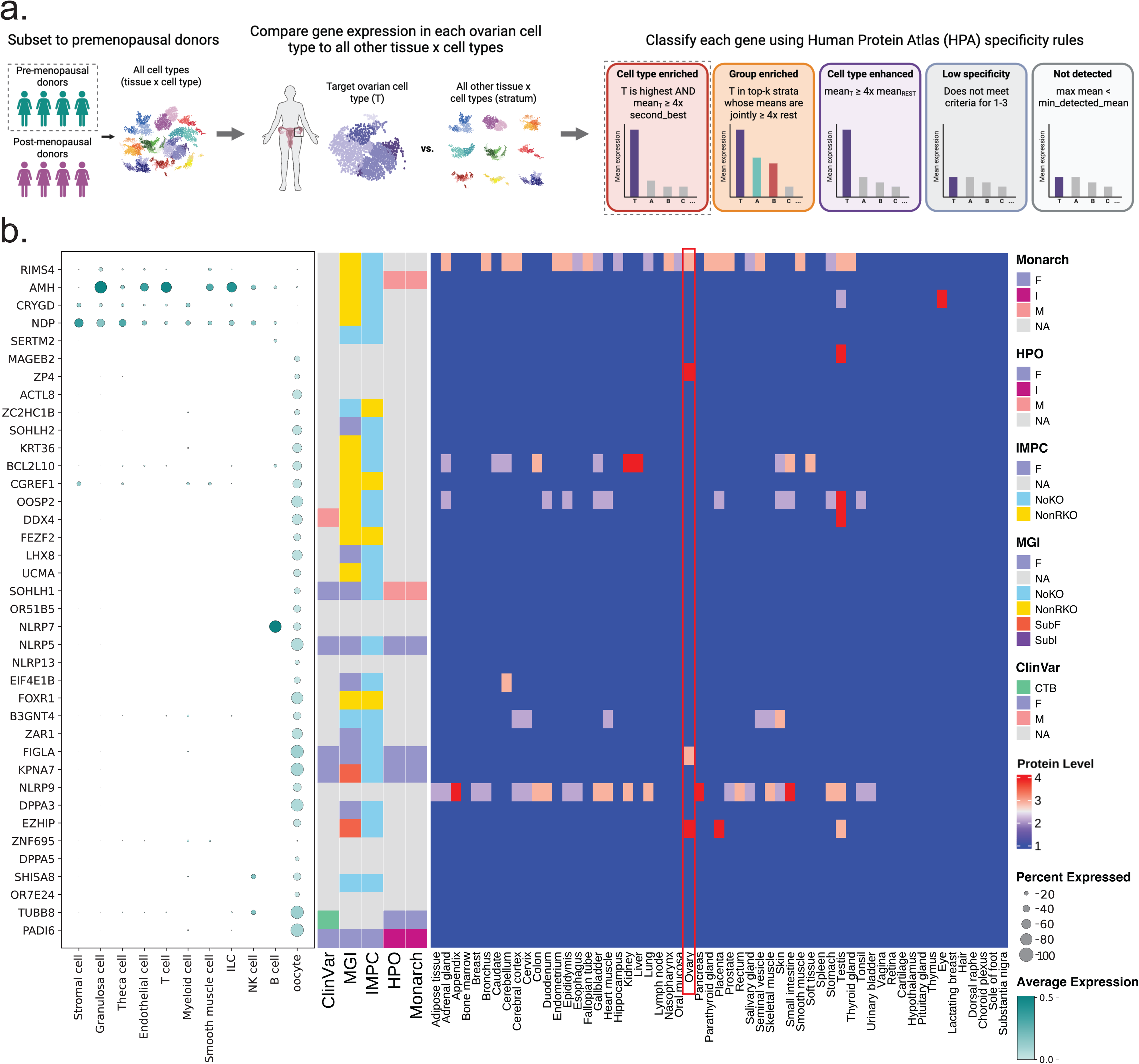
Prioritization of ovary-enriched genes using cell-type specificity and phenotype annotation databases. (a) Schematic of the ovary-specific gene prioritization workflow. Pre-menopausal donors were subset from the integrated multi-tissue atlas, and gene expression in each ovarian cell type was compared against all other ovarian and extra-ovarian cell types. Genes were classified using Human Protein Atlas-inspired specificity rules as cell type enriched, group enriched, cell type enhanced, low specificity, or not detected with respect to each ovarian cell type. Created in BioRender. VanBenschoten, H. (2026) https://BioRender.com/2xy80st (b) (Left) Dotplot and annotation heatmap of candidate ovary-enriched genes (all ovarian cell-type enriched genes). The dotplot shows average expression and percent of cells expressing each gene across each ovarian cell type. (Middle) Heatmap of mouse and human fertility phenotypes in CITDBase for each ovarian cell-type enriched gene. (Right) Protein level expression of ovarian cell marker genes across 45 tissue types in Human Protein Atlas (ref). In figure legend, CITDBase database labels are defined as follows. Monarch, HPO, and ClinVar: F = female infertility, I = non-sex specific infertility, M = male infertility, CTB = infertility in both sexes, NA = unreported. IMPC and MGI: F = female infertility, NA = unreported, NoKO = genes with no knockout, NonRKO = non-reproductive genes with a knockout, SubF = female subfertility, SubI = non-sex specific subfertility.

To further contextualize these ovarian-enriched genes, we integrated evidence from multiple phenotype and clinical annotation resources using the Contraceptive Infertility Target Database (CITDbase)^59^, including Monarch, HPO, IMPC, MGI, and ClinVar annotations (**Fig. 6b, Supplemental Data 3**). Many oocyte-enriched genes were associated with partial or full infertility phenotypes in mice and human across one or more annotation databases, although the density and type of evidence varied substantially by gene. For example, several highly oocyte-restricted genes carried human fertility- or female-associated phenotype annotations in Monarch, HPO, and ClinVar, such as *PADI6*, *FIGLA*, *SOHLH1*, and *KPNA7*, while others showed limited or absent clinical annotation despite strong ovarian specificity and evidence of infertility phenotypes at the model organism level in IMPC and MGI. This contrast highlights a key use case for the prioritization framework for distinguishing well-characterized reproductive genes from ovary-specific genes that may represent underexplored candidates.

Unsurprisingly, across the broader candidate list, specificity at the RNA level and the protein level as supported by evidence from HPA were not always concordant. Some genes with strong ovarian or oocyte-restricted expression had sparse proteomic database support, whereas a few genes, such as *EZHIP*, *FIGLA*, and *ZP4* showed medium to high protein level restricted to the ovary relative to HPA reference tissues. This may represent true biological signal in the form of translation levels and post-translation modifications or could indicate a data gap in the ovarian proteome. Overall, these results identify a set of pre-menopausal age ovary-enriched genes, particularly oocyte-associated candidates, and demonstrate how atlas-derived specificity metrics can be combined with external phenotype databases to nominate genes for downstream reproductive biology and target discovery analyses.

## DISCUSSION

In this study, we present the Menopause Cell Map, or MenoMap, a harmonized multi-tissue single-cell resource designed to examine ovarian biology in the context of reproductive aging and whole-body tissue specificity. This resource builds from high-quality existing curation and harmonization efforts, leveraging well-annotated datasets to perform a systematic comparison across cell types and tissues as a function of menopausal age. Large-scale human cell atlas efforts have transformed the study of tissue biology by enabling systematic comparison of cell types across organs and donors. However, ovarian tissue remains underrepresented in many broad atlas resources, and menopausal status is often not treated as an explicit biological variable. This is a critical limitation for reproductive biology, because the ovary undergoes profound changes across the lifespan, including follicular depletion, loss of cyclic endocrine function, stromal and vascular remodeling, and immune-state changes. By integrating more than two million cells across 13 tissues from pre- and post-menopausal age donors, the MenoMap provides a resource for distinguishing transcriptional programs associated with the functional reproductive-age ovary from those associated with the post-menopausal state.

Leveraging the MenoMap, we found that menopausal status shapes gene expression in a cell-type- and tissue-specific manner. Pre- and post-menopausal age cells broadly occupied shared transcriptional space, suggesting that major cell identities were preserved after integration and that menopausal age status did not dominate global cell-type structure. However, cell-type-resolved differential expression analyses revealed substantial menopause-associated transcriptional remodeling within specific tissue-cell-type combinations. This is consistent with prior ovarian aging studies showing that reproductive decline involves not only depletion of follicles, but also remodeling of the somatic ovarian niche, including stromal fibrosis, vascular changes, immune remodeling, oxidative stress, and cellular senescence.^46,60^ Single-cell studies of aging mouse, primate, and human ovaries have likewise identified cell-type-specific aging programs across stromal, endothelial, immune, and follicle-associated compartments, supporting the interpretation that menopausal aging is best understood at cellular resolution rather than as a uniform tissue-level state.^47,61–63^ This distinction is important as menopausal effects may be subtle or localized in global embeddings, yet highly consequential within individual cell populations. The ovary showed prominent and expected changes, especially among stromal, endothelial, immune/myeloid, and reproductive cell compartments. These findings support the broader principle that donor age and menopausal status should not be ignored during atlas-scale analysis, but can instead serve as an interpretable biological axis that influences cell state, gene expression, and downstream tissue-specificity classification.

Our ovarian-focused analyses further indicate that the post-menopausal ovary is not simply a depleted or inactive organ, but an actively remodeled tissue environment.^60^ As expected, the post-menopausal ovary showed loss of follicular cell populations and reduced representation of reproductive cell states.^55^ However, the remaining ovarian tissue was characterized by coordinated transcriptional and communication changes involving stromal, endothelial, smooth muscle, and immune populations. Cell-cell communication analysis suggested a shift from a diverse pre-menopausal network involving reproductive, stromal, vascular, and immune cell types toward a more restricted post-menopausal network dominated by somatic and vascular compartments. This was accompanied by enrichment of extracellular matrix remodeling, inflammatory signaling, vascular remodeling, stress-response, senescence, and fibrosis-associated programs. These findings are consistent with prior studies implicating stromal remodeling, inflammation, oxidative stress, and fibrosis as hallmarks of ovarian aging.^18,64,65^ The stromal and endothelial compartments emerged as particularly important sites of menopause-associated remodeling. Stromal cells showed extensive differential expression and altered ligand–receptor signaling, including pathways related to extracellular matrix organization, integrin signaling, chemokine signaling, and fibrotic remodeling. This aligns with prior work showing that ovarian aging is associated with increased stromal fibrosis and altered fibroblast, myofibroblast, and immune cell states.^66,67^ Endothelial cells also displayed substantial transcriptional changes, including signatures consistent with vascular remodeling and altered angiogenic signaling. These results are consistent with recent findings highlighting the central role of vascular and stromal niches in supporting follicle development, hormone delivery, tissue remodeling, and immune surveillance and their significant remodeling during ovarian aging. ^68,69^ Indeed, emerging evidence suggests that vascular remodeling may actively drive midlife fertility decline and the menopausal transition, positioning endothelial and stromal cells as central mediators rather than merely downstream consequences of ovarian aging.^46^ These results also suggest that post-menopausal ovarian biology should be interpreted through the state of the residual somatic microenvironment, rather than solely through the disappearance of germline and granulosa populations.

Cross-species integration with young and aged mouse ovarian single-cell datasets revealed partial conservation of ovarian programs, particularly within stromal and endothelial populations. Although several metabolic and stress-response pathways showed species- or cell-type-specific patterns, TNFα/NF-κB signaling increased consistently in postmenopausal human and aged mouse cells, suggesting that inflammatory activation represents a conserved feature of ovarian aging. Cross-species comparison of ligand–receptor interactions further identified both shared and species-specific remodeling of stromal and endothelial communication networks, underscoring the utility of mouse models for studying selected aspects of human ovarian aging while highlighting important limitations in their translational correspondence. These shared signatures align with prior single-cell studies of aging mouse, primate, and human ovaries, which have consistently identified ovarian aging as a cell-type-specific process involving stromal expansion, immune remodeling, vascular dysfunction, fibrotic extracellular matrix remodeling, and loss or alteration of germline-associated compartments.^14,18,62,64^ The convergence of these programs across species supports the interpretation that stromal and endothelial remodeling are not simply secondary consequences of follicle depletion, but may represent conserved features of the aging ovarian niche. Interestingly, we found that not all signatures were fully conserved across species or cell types. For example, steroidogenic programs showed partial divergence, particularly in endothelial cells, highlighting the biological differences between human menopause and chronological ovarian aging in mice. Human menopause reflects a distinct endocrine transition accompanied by near-complete follicular depletion, whereas mouse aging occurs in a compressed lifespan and does not fully recapitulate the timing, hormonal context, or tissue trajectory of the human menopausal transition.^63^ These distinctions are important for translational interpretation. Mouse models remain essential for mechanistic perturbation and in vivo validation, but cross-species conservation cannot be assumed for every pathway or cell state. By identifying aging-associated programs that are shared across human and mouse, as well as those that are species- or cell-type-specific, the MenoMap provides a framework for prioritizing hypotheses most likely to translate between experimental models and human ovarian biology.

Beyond differential expression and communication analysis, our results show that menopausal age can alter how tissue- and cell-type-specific gene expression is classified. Although the overall distribution of specificity categories was broadly similar between pre- and post-menopausal atlases, individual genes moved between specificity classes. The most prominent example was the loss of oocyte-associated specificity in the post-menopausal atlas, reflecting the absence of retained oocytes among post-menopausal donors in this dataset. This finding has important implications for gene specificity resources and target investigation. A gene may appear poorly expressed or non-specific in a reference dominated by post-menopausal ovarian tissue, yet be highly specific to functional reproductive-age ovarian cell populations. Conversely, some post-menopausal signals may reflect stromal, immune, or vascular remodeling rather than reproductive function. These observations underscore the need to incorporate donor life stage when interpreting ovary-enriched genes, especially for applications in fertility, reproductive disease, ovarian aging, and target discovery.

Our prioritization analysis illustrates how a multi-tissue, menopause-aware atlas can be used to nominate candidate genes for downstream biological and translational investigation. By comparing ovarian cell types against both ovarian and extra-ovarian populations, we recovered expected oocyte-enriched genes, including *NLRP5*, *NLRP14*, *FIGLA*, *KPNA7*, *NLRP9*, *PADI6*, *ZP4*, and *OOSP2*. Many of these genes have prior evidence linking them to oocyte biology, fertility, or reproductive phenotypes, while others remain less well characterized in human systems. Integrating RNA-level specificity with Human Protein Atlas protein evidence and fertility phenotype annotations provides a more nuanced prioritization framework than expression specificity alone. This is particularly important because RNA expression does not necessarily imply protein abundance, functional requirement, druggability, or safety as a therapeutic target.^11,70,71^ Indeed, in the nonhormonal contraception use-case many of the genes in the cell type enriched list are not ideal targets due to their involvement in development processes, intracellular location, or potential non-reversibility. Thus, rather than defining a final list of targets, this atlas provides a structured approach for triaging ovary-enriched genes according to cell-type specificity, menopausal context, protein-level support, and genetic or model-organism evidence.

We would also like to temper interpretations with a few limitations of our work. Menopausal status is also closely intertwined with chronological age and other donor-level variables, including hormonal history, parity, medication use, and overall health status, which are not uniformly available across public datasets.^36^ Because the data are cross-sectional, we cannot distinguish causal effects of menopause from correlated effects of aging or other unmeasured variables. Additionally, menopause occurs at different ages across individuals and is marked by a perimenopausal period of up to 5 years which is not included in the metadata of this resource.^37,38^ Future work should aim to incorporate this peri-menopausal transition and very clear markers of menopause such as hormone levels into analyses. Single-cell curation efforts should also seek to obtain explicit information about menopausal status in female donors where possible. Finally, atlas construction from public datasets is inherently limited by data availability.

Despite these limitations, the MenoMap provides a comprehensive resource for studying ovarian biology in a whole-body and lifespan-aware context. By explicitly incorporating pre- and post-menopausal donors, the atlas helps separate features of the functional reproductive-age ovary from those associated with post-menopausal remodeling and more broadly understand the transcriptomic shifts across tissues. This framework can support studies of ovarian aging, reproductive disease, fertility biology, and candidate target investigation, while also providing a model for how life-stage variables can be incorporated into human cell atlas resources and drug discovery efforts. Future work integrating spatial transcriptomics, proteomics, longitudinal sampling, and experimentally validated perturbation data will further refine the biological and translational utility of this resource. This study also highlights that donor metadata such as menopausal status, age, tissue source, and health status can significantly affect biological interpretation and should be curated and modeled wherever possible. Finally, we find that tissue specificity is not an immutable property of a gene, but a context-dependent classification that can change with cell composition and donor life stage. This is especially relevant for reproductive tissues, where the presence or absence of transient, rare, or life-stage-specific cell populations can strongly influence apparent tissue enrichment.

## MATERIALS AND METHODS

### Dataset curation

We curated scRNA-seq datasets from CZ CellXGene that met inclusion criteria for donor metadata and sequencing modality.^72^ We chose CZ CellXGene because this is a harmonized resource that provides robust access to published datasets that are well-suited for our work and enable downstream processing steps that reflect the original published datasets (see next section). Specifically, we included datasets containing human female developmentally mature (Adult – 15+ years) donors and included only healthy tissue samples (Disease status = normal). Data were further filtered by assay to include only samples sequenced via 10x 3’ v1-v3 methods. One additional study external to CellXGene was curated and preprocessed to incorporate age-specified oocytes^14^, resulting in a total of 20 individual datasets encompassing 13 tissue types and 278 unique female donors. Upon application of this filtering criteria, the 13 tissue types included as references were adipose, breast, eye, fallopian tube, heart, intestine, kidney, liver, lung, ovary, pancreas, skin, and uterus. Any gaps in tissue type reflect the lack of data that met the specified inclusion criteria from the CellXGene database at the time of data curation (November 2025). All donor and study-level metadata are recorded in **Supplemental Data 1**.

### Dataset preprocessing and cell type harmonization

Cell type nomenclature was harmonized by semantic hierarchical cell type harmonization for each tissue type using the CellXGene-provided cell_type column.^72^ All CellXGene and author-defined cell type mappings were retained in the final object. Detailed mappings of harmonized cell type labels are provided in **Supplemental Data 2**. The Scanpy environment was used to load preprocessed h5ad files into AnnData objects and perform the following quality control procedures.^75^ First, Scrublet was applied to each individual sample to exclude doublets and ensure that only high-quality singlets were maintained in the final dataset.^76^ For each dataset, cells were retained if they expressed more than 400 genes and had less than 20% mitochondrial and ribosomal gene content; genes were retained if detected in more than three cells. All datasets except for breast had <500,000 cells, thus only breast underwent density-aware down sampling to maintain order-of-magnitude similarity in cell number across tissues. A downsampling procedure was applied to oversized datasets to reduce their size to <500,000 cells using density-aware subsampling. All cells from rare annotated types (<0.1% of cells or <100 cells per type) were kept; common types were subsampled using probabilities inversely related to local density in PCA space (15-neighbor mean distance, computed in 50,000-cell chunks), so dense regions were downsampled more aggressively than sparse ones. Sampling was reproducible (seed = 42). Lastly, the gene_biotype column was used to identify protein-coding genes, and only protein coding genes were used for integration and downstream analysis (19,400 genes total).

For the manually curated and preprocessed oocyte-containing ovary dataset, data was retrieved from GSE255690^14^ and a standard Seurat pipeline was performed to annotate and subset high-confidence oocytes and granulosa cells.^77^ Briefly, Single-cell RNA-seq datasets from 9 ovarian donors spanning young, middle-aged, and old groups were imported into Seurat, merged into a unified object, and annotated with donor-level metadata. Cells with >20% mitochondrial or ribosomal transcript content were removed prior to normalization, identification of 2,000 highly variable genes, scaling with regression of mitochondrial and ribosomal percentages, PCA dimensionality reduction, graph-based clustering using the top 20 principal components, and UMAP visualization; cluster stability was evaluated across resolutions from 0.2–0.5, with a final clustering resolution of 0.2 selected for downstream analyses. Cell identities were assigned using canonical marker expression and module scoring against reference-derived signatures, and refined oocyte and granulosa populations were subsetted and exported as an .h5ad object for downstream analysis.

Prior to atlas integration, we harmonized AnnData structure across sources. CELLxGENE objects store library-size-normalized, log-transformed expression in adata.X with unnormalized counts in .raw and Ensembl gene identifiers. For non-CELLxGENE datasets, we copied raw counts before normalization, applied library-size normalization (10,000 counts per cell) and log1p transformation to adata.X, mapped gene symbols to Ensembl IDs (MyGene; genes without mappings were excluded), stored the original count matrix in adata.raw, added harmonized sample metadata (tissue, tissue_id), and standardized sparse matrix format (CSR, float32) while removing auxiliary layers. This ensured a consistent representation of normalized expression and raw counts across datasets prior to merging.

### Atlas integration and benchmarking

Raw count layers of each QC-processed dataset were merged using sc.concat with unique cell indices and union of all genes. scVI and scANVI were used to generate a donor-corrected latent embedding of the full atlas according to the following method and parameters.^34,35^ First, scVI was fit on raw counts with batch correction by unique donor id, using a 2-layer encoder/decoder, 30 latent dimensions, and a negative binomial gene likelihood. scVI was trained for 400 epochs (validation every 50 epochs; default scVI learning rate ∼1×10⁻³). The scVI latent representation was stored as X_scVI. Then, scANVI was initialized from the trained scVI weights and trained on harmonized cell-type labels (h_cell_type, with Unknown as the unlabeled category) for 20 epochs with 100 cells sampled per label per epoch and learning rate 5×10⁻ (reduced from the default to improve stability with large gene panels). Convergence was confirmed via stability in the Evidence Lower Bound (ELBO) loss curve. The final latent embedding was saved as X_h_scANVI in an integrated .h5ad file. Random seeds were fixed (seed = 0). Scib-metrics was used to benchmark X_scANVI, X_scVI, and unintegrated X_PCA embeddings with respect to bio conservation and batch correction metrics.^78^

### Pre- and post-menopausal donor labelling

Premenopausal and postmenopausal labels were inferred from development_stage metadata using hierarchical text rules (**Table 1**) and stored as menopausal_status on the integrated atlas.

| Order | Condition (case-insensitive substring / pattern) | Label |
| --- | --- | --- |
| — | Missing, empty, nan, or none | unknown |
| 1 | Contains young adult | premenopausal |
| 2 | Contains third decade **or** fourth decade | premenopausal |
| 3 | Matches N-year-old, N year old, or N years old | premenopausal if $N \leq 50$; postmenopausal if $N > 50$ |
| 4 | Contains adult stage | premenopausal |
| 5 | Contains decade **or** postnatal (and not already matched above) | postmenopausal |
| 6 | First integer in string | premenopausal if $\leq 50$; postmenopausal if $> 50$ |

### Marker gene and differential gene expression analysis

To identify tissue marker genes (Fig. 1e), differential expression was computed with the trained scANVI model (mode=“change”) comparing each tissue to all others on raw counts (counts layer), with batch correction by donor–tissue using the scvi-tools differential_expression() function. Upregulated genes per tissue were ranked by posterior mean log-fold change (lfc_mean); genes were filtered by FDR < 0.05, and the top five genes per tissue were reported. Dot plots used library-size-normalized, log-transformed counts for the selected marker genes.

To identify differentially expressed genes by menopause status for each tissue x cell type without treating cells as independent biological replicates, we performed donor-level pseudobulk differential expression. For each tissue and h_cell_type with ≥15 premenopausal and ≥15 postmenopausal cells, raw counts (counts layer) were summed per donor (donor_id) using decoupler pseudobulking (mode=’sum’). Pseudobulk samples with <10 cells or <1,000 total counts were excluded. Menopausal status was assigned at the donor level (majority label among that donor’s cells in the stratum), and only donors labeled premenopausal or postmenopausal were retained. Strata with fewer than two donors in either group were skipped.

Lowly expressed genes were filtered with decoupler (filter_by_expr / filter_by_prop). Differential expression was then tested with pyDESeq2^79^ using the design ∼ menopausal_status, with premenopausal as the reference level and the contrast postmenopausal vs premenopausal. Genes with Benjamini–Hochberg adjusted *P* < 0.05 and |log₂ fold change| > 1 were counted as significant. Counts of significant DEGs were summarized as a tissue × cell-type heatmap.

Differential expression among ovarian cell types between menopausal states (postmenopausal vs. premenopausal in human; old vs. young in mouse) was performed independently for each major cell type using a pseudobulk approach or a mixed-model framework implemented within Seurat, as appropriate for the dataset structure.^77^ Genes were considered significantly differentially expressed at an adjusted p-value threshold of 0.05 (Benjamini–Hochberg correction) and an absolute log₂ fold-change threshold of 0.25. Upward- and downward-regulated gene sets were retained separately for downstream enrichment and visualization analyses.

### Differential cell-type proportion analysis and compositional abundance testing

For each tissue with both premenopausal and postmenopausal cells, we computed the proportion of each harmonized cell type within each menopause group. Post/pre fold changes were calculated as FC = (prop_post + ε) / (prop_pre + ε), with pseudocount ε = 10⁻. For cell types present in only one menopause group within a tissue, FC was capped to the maximum bilateral FC observed in that tissue (or 2.0 if no bilateral types were available), assigned as FC = M for post-only types and FC = 1/M for pre-only types. Every cell type observed in either menopause group within a tissue was included in the resulting tissue × cell-type matrix. To identify menopause-associated compositional changes while accounting for donor-to-donor variation, we applied scCODA separately within each tissue.^44^ For each tissue, cell-type count tables were constructed at the sample level using donor id as the replicate unit. Each sample was assigned a single menopausal status (modal status across its cells). Tissues were analyzed only if they contained at least two menopause groups with ≥2 samples per group and at least two cell types with nonzero counts. We fit scCODA compositional models (CompositionalAnalysis) with formula menopausal_status, automatic reference cell-type selection, and Hamiltonian Monte Carlo inference (8,000 posterior samples after 2,000 burn-in iterations). Credible effects were defined using scCODA’s inclusion-based FDR calibration at a target FDR of 0.1. A tissue-cell-type pair was marked as significant if that cell type showed a credible menopause effect in the scCODA model run for that tissue.

### Subsetted human and mouse ovary single-cell RNA-sequencing data processing

Human ovarian scRNA-seq data were subset from the integrated AnnData object using the tissue column (tissue = ovary) and downstream analysis was performed using Seurat (v5).^77^ Raw count matrices were log-normalized, scaled to 10,000 counts per cell, and variance-stabilized. Highly variable features were identified using the vst method, and dimensionality was reduced by principal component analysis (PCA) followed by uniform manifold approximation and projection (UMAP) for visualization. Cells were clustered using the Leiden algorithm at empirically determined resolutions and annotated to major cell types based on canonical marker gene expression. The final annotated object retained cells with a valid cell-type label (h_cell_type) and a defined menopausal status (premenopausal or postmenopausal). Mouse scRNA-seq datasets were processed with an analogous workflow. Where UMAP embeddings were absent, the standard pipeline (NormalizeData, FindVariableFeatures (n = 2,000), ScaleData, RunPCA (30 PCs), RunUMAP, FindNeighbors, FindClusters) was applied. For Seurat v5 objects, split layers were merged with JoinLayers prior to downstream analysis. Cell-type labels from the Pepin^80^ & Goods^27^ mouse dataset were harmonized to a common vocabulary using a curated mapping table, consolidating granulosa sub-states, epithelial, pericyte, smooth muscle and immune populations into defined categories.

### Ovarian gene set enrichment and overlap analysis

Curated gene sets representing biological programs of established relevance to ovarian physiology, vascular biology and reproductive ageing were compiled from the literature and public databases. These encompassed processes including ECM remodeling and fibrosis, endothelial identity and barrier function, steroidogenesis, cellular senescence and the senescence-associated secretory phenotype (SASP), angiogenesis and lymphangiogenesis, TGF-β signaling, estrogen and progesterone response, IL-6/JAK/STAT3 signaling, TNFα/NFκB signaling, hypoxia response, ovarian reserve and folliculogenesis, and endometriosis-associated stromal programs, among others. For each gene set and each cell type, the overlap with the significant DEG list was quantified. Enrichment significance was evaluated by a one-sided hypergeometric test using the full tested gene universe as background and resulting p-values were adjusted by the Benjamini–Hochberg procedure. Results were visualized as waffle-tile maps in which each tile represents a single DEG, colored by direction of regulation.

### Inference of cell-cell communication

Ligand–receptor-mediated intercellular communication was inferred using CellChat (v2) applied to each condition independently.^81^ Normalized expression data and cell-type metadata were passed to createCellChat, with the curated CellChatDB.human or CellChatDB.mouse interaction database as appropriate. Over-expressed signaling genes and interactions were identified with identifyOverExpressedGenes and identifyOverExpressedInteractions. Communication probabilities were computed with computeCommunProb, and cell-type populations contributing fewer than ten cells were removed by filterCommunication. Pathway-level probabilities were then aggregated with computeCommunProbPathway, and network-level centrality metrics were derived with netAnalysis_computeCentrality. Differential interaction analysis between conditions was based on log₂-transformed fold-changes in mean communication probability for each ligand–receptor pair and sender–receiver cell-type combination.

### Cross-species integration and module scoring

To assess the evolutionary conservation of menopausal transcriptional programs, human and mouse ovarian datasets were jointly embedded using Seurat’s anchor-based integration framework. Human genes were mapped to their corresponding mouse orthologues using Ensembl BioMart, and only genes with identified cross-species orthologues were retained for the integration and comparative analyses. Premenopausal human cells were aligned with young mouse cells, whereas postmenopausal human cells were aligned with old mouse cells. Integration anchors were identified with FindIntegrationAnchors and harmonized embeddings were computed with IntegrateData, followed by PCA and UMAP. Gene module scores for steroidogenesis, cellular senescence, oxidative stress and TGF-β/fibrosis programs were computed with AddModuleScore using the curated gene sets described above. Score distributions were compared across conditions and species with the Wilcoxon rank-sum test; p-values were adjusted for multiple comparisons. Conservation of differential LR interactions between the human and mouse systems was quantified by Pearson correlation of the respective log₂FC vectors, restricted to interaction pairs detected in both species.

### Atlas-wide and ovary specific cell type specificity characterization

We classified each gene’s specificity among cell types using HPA-inspired rules on stratum-averaged expression (scANVI batch-corrected normalized expression, library size 10, log-transformed; means per tissue×harmonized cell type).^22^ Genes were labeled as not detected, low specificity, cell type enriched (top stratum ≥4-fold above second), group enriched (top 2–10 strata jointly ≥4-fold above remainder), or cell type enhanced (≥4-fold above mean of others). Labels were computed separately for premenopausal and postmenopausal cells (menopausal status from metadata or age ≥50). Pre- to post-menopausal transitions between cell type specificity categories per gene were visualized as an Alluvial flow diagram (Fig. 4b); gains and losses in the three high-specificity tiers were summarized by tissue and cell type and plotted as barplots (Fig. 4c&d). Over-representation of gained and lost gene sets (union across tiers) was tested with g:Profiler against GO (biological process, molecular function, cellular component), with Benjamini–Hochberg adjustment; additional GO ORA was run for selected tissue-specific contrasts (Fig. 4e&f). Using the same fold and detection parameters, we classified genes with respect to each premenopausal ovarian cell type against all premenopausal cell-type means atlas-wide, counted genes per specificity category, and exported cell-type–enriched gene lists. Selected genes were visualized in dot plots of expression across premenopausal ovarian cell types (library-size normalized, log-transformed counts) (Fig. 6b).

### Visualization

Visualizations containing graphical abstracts or workflows (Fig. 1a&b, Fig. 5a, Fig. 6a) were generated using BioRender with licenses in respective figure panel captions. All visualizations for Fig. 1, 2, 5 and 6 were generated using matplotlib in Python. UMAPs were computed with Scanpy and drawn with Scanpy plotters sc.pl.umap or sc.pl.embedding; dotplots for marker genes used Scanpy sc.pl.dotplot; and heatmaps used matplotlib imshow and seaborn heatmap. All visualizations for Fig. 3 and 4 were generated in R using ggplot2, patchwork, circlize, ggalluvial, ComplexHeatmap, UpSetR and CellChat plotting utilities. Chord diagrams depicting global communication networks were produced with the circlize package. Alluvial diagrams of sender–receiver interaction flow were generated with ggalluvial. UpSet plots were drawn with UpSetR. Lollipop and dot-plot panels were implemented with custom ggplot2 geoms. Waffle-tile enrichment maps used geom_tile with per-gene directional coloring. All figures were assembled at a uniform base font size of 9 pt (Helvetica/Arial) following Nature-portfolio style guidelines and exported as vector PDF via cairo_pdf or the pdf device.

### Statistical analysis

Unless otherwise stated, multiple-testing correction was applied using the Benjamini–Hochberg false discovery rate procedure. Significance thresholds are denoted as follows: *, adjusted p < 0.05; **, adjusted p < 0.01; ***, adjusted p < 0.001; ns, not significant. Sample sizes (cell numbers per condition and per cell type) are reported in the corresponding figure panels and supplementary tables.

## ACKNOWLEDGEMENTS

We would like to thank Dr. Margie Ackerman Lab and Dr. Jiwon Lee, and their lab members for helpful discussions throughout this project. We would also like to thank Dr. Shuo Xiao, Dr. Jeff Pea, and Dr. Stephen Ward for helpful discussions. This work was supported by the Bill & Melinda Gates Foundation [INV-003385 and INV-040475]. Under the grant conditions of the Foundation, a Creative Commons Attribution 4.0 Generic License has already been assigned to the Author Accepted Manuscript version that might arise from this submission. B.A.G is supported in part through the Geisel School of Medicine at Dartmouth’s Center for Quantitative Biology through a grant from the National Institute of General Medical Sciences (NIGMS, P20GM130454) of the NIH.

## DATA AVAILABILITY

Source data are available in the public datasets and databases in **Supplemental Data 1**. The integrated Menopause Cell Map is available via: https://cellinteractionsdb.shinyapps.io/shinyapp_premeno_scrna-seq/.

## CODE AVAILABILITY

All code for MenoMap assembly is available on GitHub at: github.com/Goods-Lab/MenoMap-Integration.git

