## Supplemental Figures for "A single-cell atlas inclusive of age-related menopause reveals global tissue and ovarian remodeling"

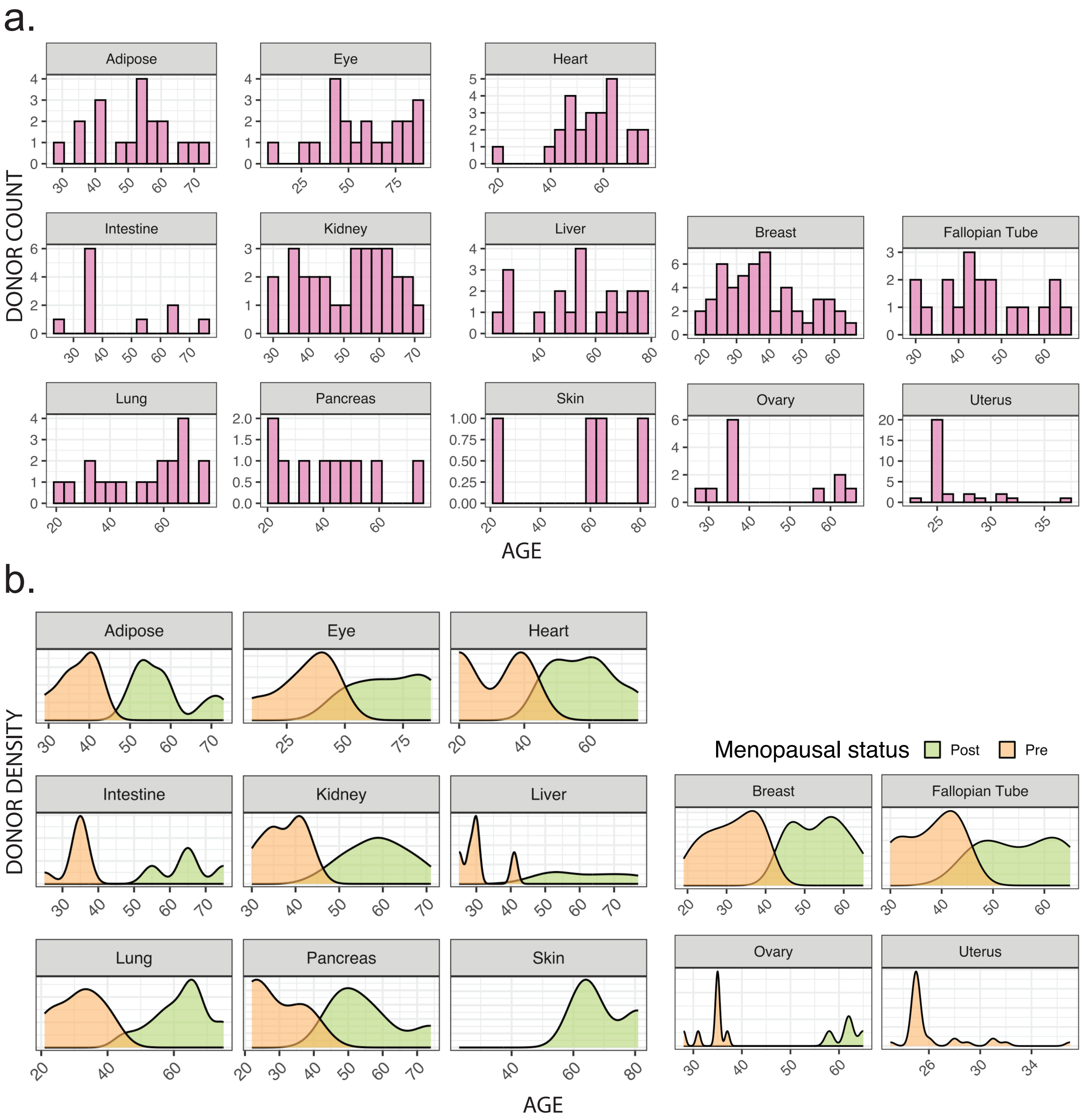


**Supplemental Figure 1. Distribution of donor age and menopause status per tissue**

**(a)** Histogram of donor age per tissue in MenoMap.

**(b)** Density plot of donor menopause status as a function of age per tissue in MenoMap. Orange correspond to premenopausal; green corresponds to postmenopausal.

| Method | Bio conservation | | | | Batch correction | | | | | Aggregate score | | |
|---|---|---|---|---|---|---|---|---|---|---|---|---|
| | KMeans NMI | KMeans ARI | Silhouette label | cLISI | BRAS | iLISI | KBET | Graph connectivity | PCR comparison | Batch correction | Bio conservation | Total |
| **h_scANVI** | 0.61 | 0.21 | 0.48 | 1.00 | 0.65 | 0.01 | 0.46 | 0.91 | 0.63 | 0.53 | 0.58 | 0.56 |
| **Unintegrated** | 0.71 | 0.29 | 0.50 | 1.00 | 0.50 | 0.00 | 0.30 | 0.67 | 0.00 | 0.29 | 0.62 | 0.49 |

**Supplemental Figure 2. scANVI integration benchmark assessment.**

Scib-metrics table of scANVI integration compared to unintegrated PCA embedding. Numeric labels indicate absolute metrics; purple dots represent lower metric values; green dots represent higher metric values.

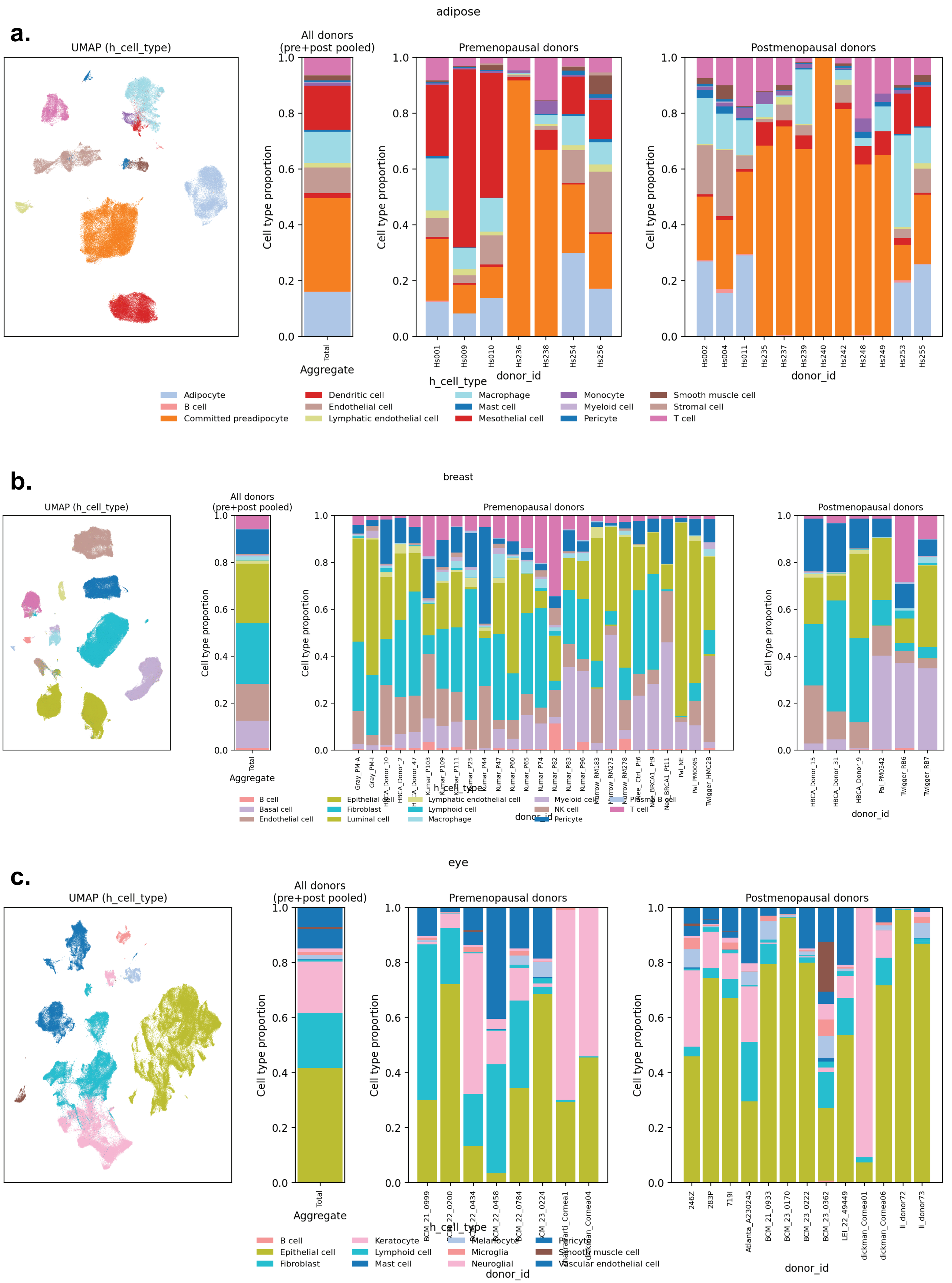
a.
adipose
UMAP (h_cell_type)
All donors
(pre+post pooled)
Premenopausal donors
Postmenopausal donors
Cell type proportion
Total
Aggregate
Hs001
Hs009
Hs010
Hs236
Hs238
Hs254
Hs256
Hs002
Hs004
Hs011
Hs235
Hs237
Hs239
Hs240
Hs242
Hs248
Hs249
Hs253
Hs255
donor_id
h_cell_type
Adipocyte
B cell
Committed preadipocyte
Dendritic cell
Endothelial cell
Lymphatic endothelial cell
Macrophage
Mast cell
Mesothelial cell
Monocyte
Myeloid cell
Pericyte
Smooth muscle cell
Stromal cell
T cell
b.
breast
UMAP (h_cell_type)
All donors
(pre+post pooled)
Premenopausal donors
Postmenopausal donors
Cell type proportion
Total
Aggregate
Gray_PM-A
Gray_PM-I
Pal_NE
Pal_PM0095
Twigger_HMC2B
HBCA_Donor_15
HBCA_Donor_31
HBCA_Donor_9
Pal_PM0342
Twigger_RB6
Twigger_RB7
donor_id
h_cell_type
B cell
Basal cell
Endothelial cell
Epithelial cell
Fibroblast
Luminal cell
Lymphatic endothelial cell
Lymphoid cell
Macrophage
Myeloid cell
NK cell
Pericyte
Plasma B cell
T cell
c.
eye
UMAP (h_cell_type)
All donors
(pre+post pooled)
Premenopausal donors
Postmenopausal donors
Cell type proportion
Total
Aggregate
246Z
283P
719I
Atlanta_A230245
BCM_21_0933
BCM_23_0170
BCM_23_0222
BCM_23_0362
LEI_22_49449
dickman_Cornea01
dickman_Cornea06
li_donor72
li_donor73
donor_id
h_cell_type
B cell
Epithelial cell
Fibroblast
Keratocyte
Lymphoid cell
Mast cell
Melanocyte
Microglia
Neuroglia
Pericyte
Smooth muscle cell
Vascular endothelial cell

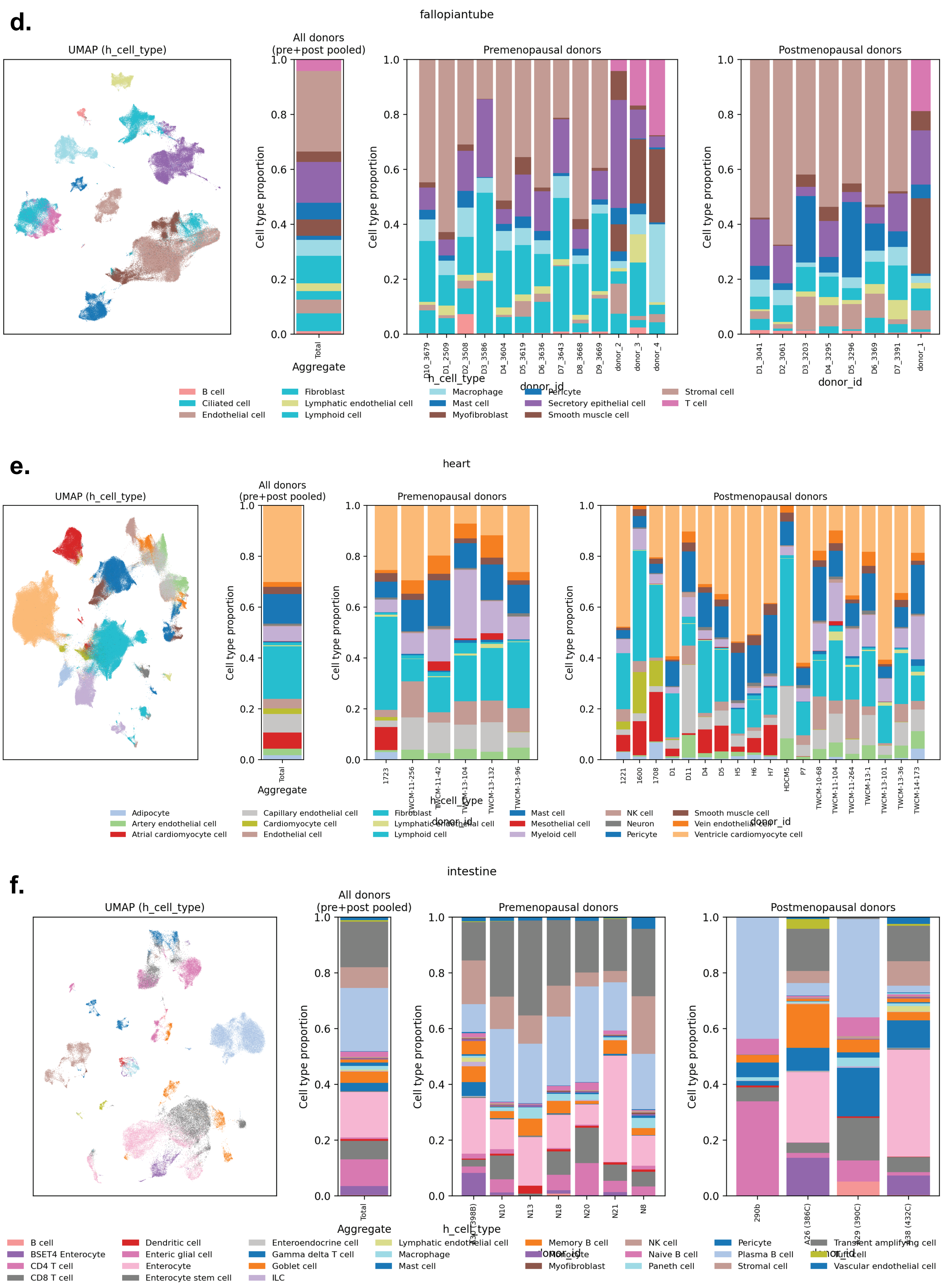
d.
fallopiantube
UMAP (h_cell_type)
All donors (pre+post pooled)
Premenopausal donors
Postmenopausal donors
Cell type proportion
Aggregate
Total
h_cell_type
donor_id
D10_3679
D1_2509
D2_3508
D3_3586
D4_3604
D5_3619
D6_3636
D7_3643
D8_3668
D9_3669
donor_2
donor_3
donor_4
D1_3041
D2_3061
D3_3203
D4_3295
D5_3296
D6_3369
D7_3391
donor_1
B cell
Ciliated cell
Endothelial cell
Fibroblast
Lymphatic endothelial cell
Lymphoid cell
Macrophage
Mast cell
Myofibroblast
Pericyte
Secretory epithelial cell
Smooth muscle cell
Stromal cell
T cell
e.
heart
UMAP (h_cell_type)
All donors (pre+post pooled)
Premenopausal donors
Postmenopausal donors
Cell type proportion
Aggregate
Total
h_cell_type
donor_id
1723
TWCM-11-256
TWCM-11-42
TWCM-13-104
TWCM-13-132
TWCM-13-96
1221
1600
1708
D1
D11
D4
D5
H5
H6
H7
HDCM5
P7
TWCM-10-68
TWCM-11-104
TWCM-11-264
TWCM-13-1
TWCM-13-101
TWCM-13-36
TWCM-14-173
Adipocyte
Artery endothelial cell
Atrial cardiomyocyte cell
Capillary endothelial cell
Cardiomyocyte cell
Endothelial cell
Fibroblast
Lymphatic endothelial cell
Lymphoid cell
Mast cell
Mesothelial cell
Myeloid cell
NK cell
Neuron
Pericyte
Smooth muscle cell
Vein endothelial cell
Ventricle cardiomyocyte cell
f.
intestine
UMAP (h_cell_type)
All donors (pre+post pooled)
Premenopausal donors
Postmenopausal donors
Cell type proportion
Aggregate
Total
h_cell_type
donor_id
N10
N13
N18
N20
N21
N8
290b
B cell
BSET4 Enterocyte
CD4 T cell
CD8 T cell
Dendritic cell
Enteric glial cell
Enterocyte
Enterocyte stem cell
Enteroendocrine cell
Gamma delta T cell
Goblet cell
ILC
Lymphatic endothelial cell
Macrophage
Mast cell
Memory B cell
Myofibroblast
NK cell
Naive B cell
Paneth cell
Pericyte
Plasma B cell
Stromal cell
Transient amplifying cell
Vascular endothelial cell

g.

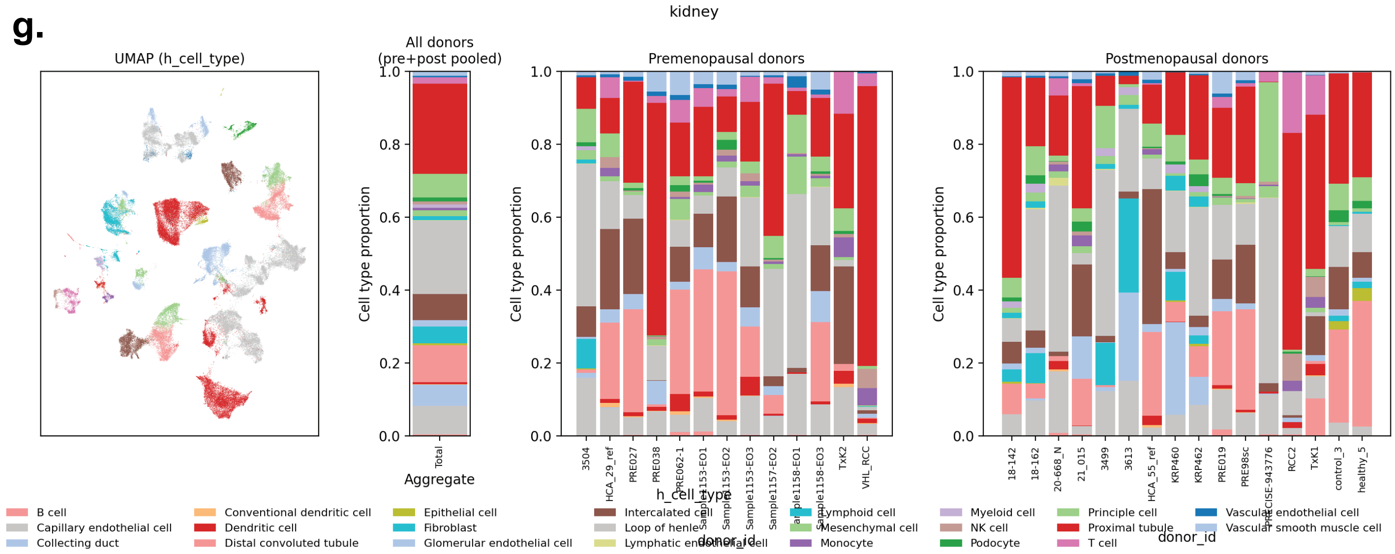

kidney
UMAP (h_cell_type)
All donors (pre+post pooled)
Premenopausal donors
Postmenopausal donors
Cell type proportion
Total
Aggregate
h_cell_type
donor_id
3504
HCA_29_ref
PRE027
PRE038
PRE062-1
Sample1153-EO1
Sample1153-EO2
Sample1153-EO3
Sample1157-EO2
Sample1158-EO1
Sample1158-EO3
TxK2
VHL_RCC
18-142
18-162
20-668_N
21_015
3499
3613
HCA_55_ref
KRP460
KRP462
PRE019
PRE98sc
PRECISE-943776
RCC2
TxK1
control_3
healthy_5
B cell
Capillary endothelial cell
Collecting duct
Conventional dendritic cell
Dendritic cell
Distal convoluted tubule
Epithelial cell
Fibroblast
Glomerular endothelial cell
Intercalated cell
Loop of henle
Lymphatic endothelial cell
Lymphoid cell
Mesenchymal cell
Monocyte
Myeloid cell
NK cell
Podocyte
Principle cell
Proximal tubule
T cell
Vascular endothelial cell
Vascular smooth muscle cell


h.

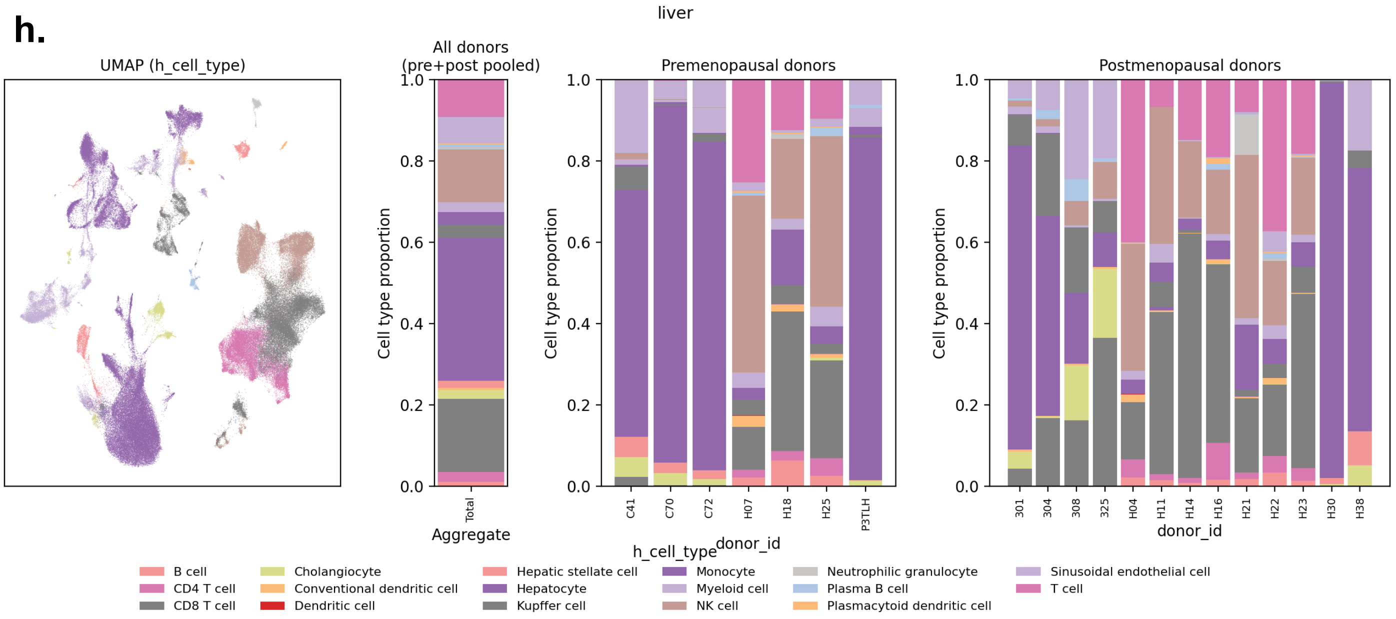

liver
UMAP (h_cell_type)
All donors (pre+post pooled)
Premenopausal donors
Postmenopausal donors
Cell type proportion
Total
Aggregate
h_cell_type
donor_id
C41
C70
C72
H07
H18
H25
P3TLH
301
304
308
325
H04
H11
H14
H16
H21
H22
H23
H30
H38
B cell
CD4 T cell
CD8 T cell
Cholangiocyte
Conventional dendritic cell
Dendritic cell
Hepatic stellate cell
Hepatocyte
Kupffer cell
Monocyte
Myeloid cell
NK cell
Neutrophilic granulocyte
Plasma B cell
Plasmacytoid dendritic cell
Sinusoidal endothelial cell
T cell


i.

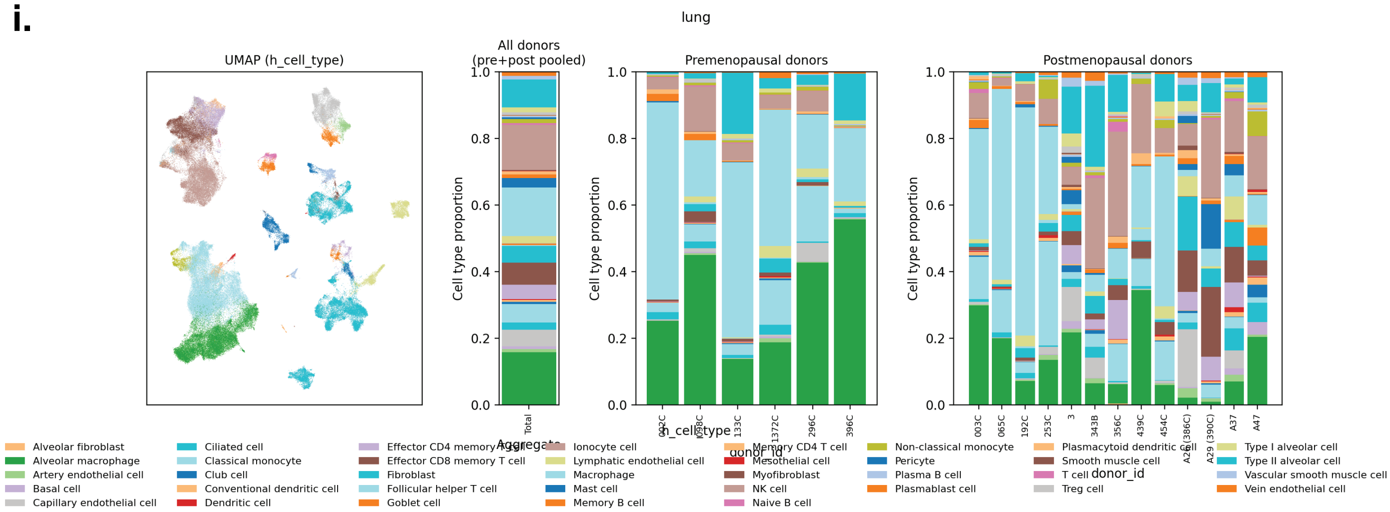

lung
UMAP (h_cell_type)
All donors (pre+post pooled)
Premenopausal donors
Postmenopausal donors
Cell type proportion
Total
Aggregate
h_cell_type
donor_id
133C
1372C
296C
396C
003C
065C
192C
253C
3
343B
356C
439C
454C
A37
A47
Alveolar fibroblast
Alveolar macrophage
Artery endothelial cell
Basal cell
Capillary endothelial cell
Ciliated cell
Classical monocyte
Club cell
Conventional dendritic cell
Dendritic cell
Effector CD4 memory T cell
Effector CD8 memory T cell
Fibroblast
Follicular helper T cell
Goblet cell
Ionocyte cell
Lymphatic endothelial cell
Macrophage
Mast cell
Memory B cell
Memory CD4 T cell
Mesothelial cell
Myofibroblast
NK cell
Naive B cell
Non-classical monocyte
Pericyte
Plasma B cell
Plasmablast cell
Plasmacytoid dendritic cell
Smooth muscle cell
T cell
Treg cell
Type I alveolar cell
Type II alveolar cell
Vascular smooth muscle cell
Vein endothelial cell

j.

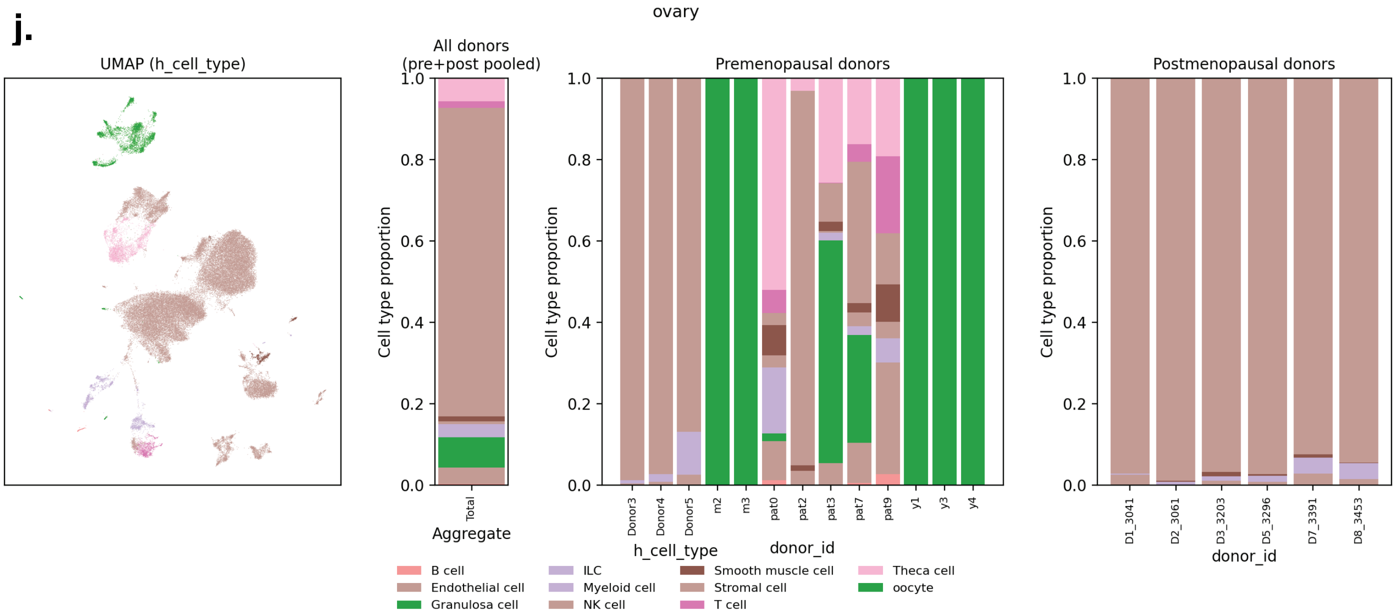
ovary
UMAP (h_cell_type)
All donors
(pre+post pooled)
Premenopausal donors
Postmenopausal donors
Cell type proportion
0.0
0.2
0.4
0.6
0.8
1.0
Total
Aggregate
Donor3
Donor4
Donor5
m2
m3
pat0
pat2
pat3
pat7
pat9
y1
y3
y4
donor_id
D1_3041
D2_3061
D3_3203
D5_3296
D7_3391
D8_3453
donor_id
h_cell_type
B cell
Endothelial cell
Granulosa cell
ILC
Myeloid cell
NK cell
Smooth muscle cell
Stromal cell
T cell
Theca cell
oocyte

k.

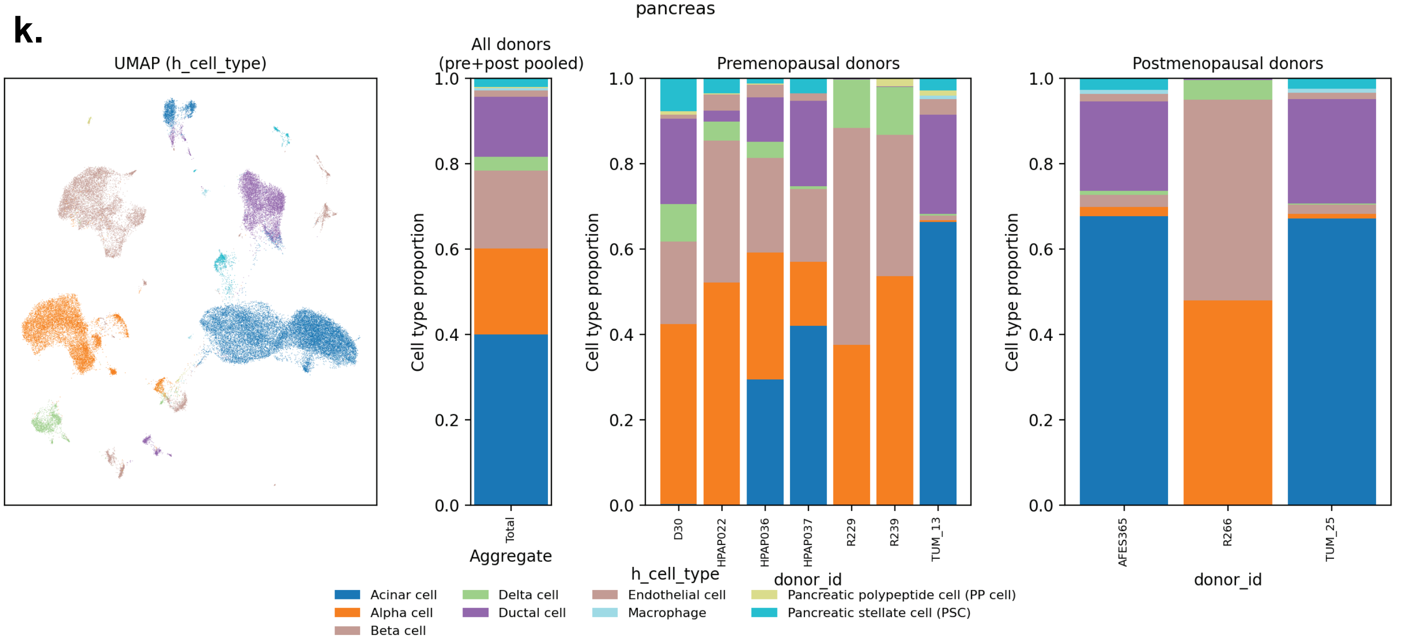
pancreas
UMAP (h_cell_type)
All donors
(pre+post pooled)
Premenopausal donors
Postmenopausal donors
Cell type proportion
0.0
0.2
0.4
0.6
0.8
1.0
Total
Aggregate
D30
HPAP022
HPAP036
HPAP037
R229
R239
TUM_13
donor_id
AFES365
R266
TUM_25
donor_id
h_cell_type
Acinar cell
Alpha cell
Beta cell
Delta cell
Ductal cell
Endothelial cell
Macrophage
Pancreatic polypeptide cell (PP cell)
Pancreatic stellate cell (PSC)

l.

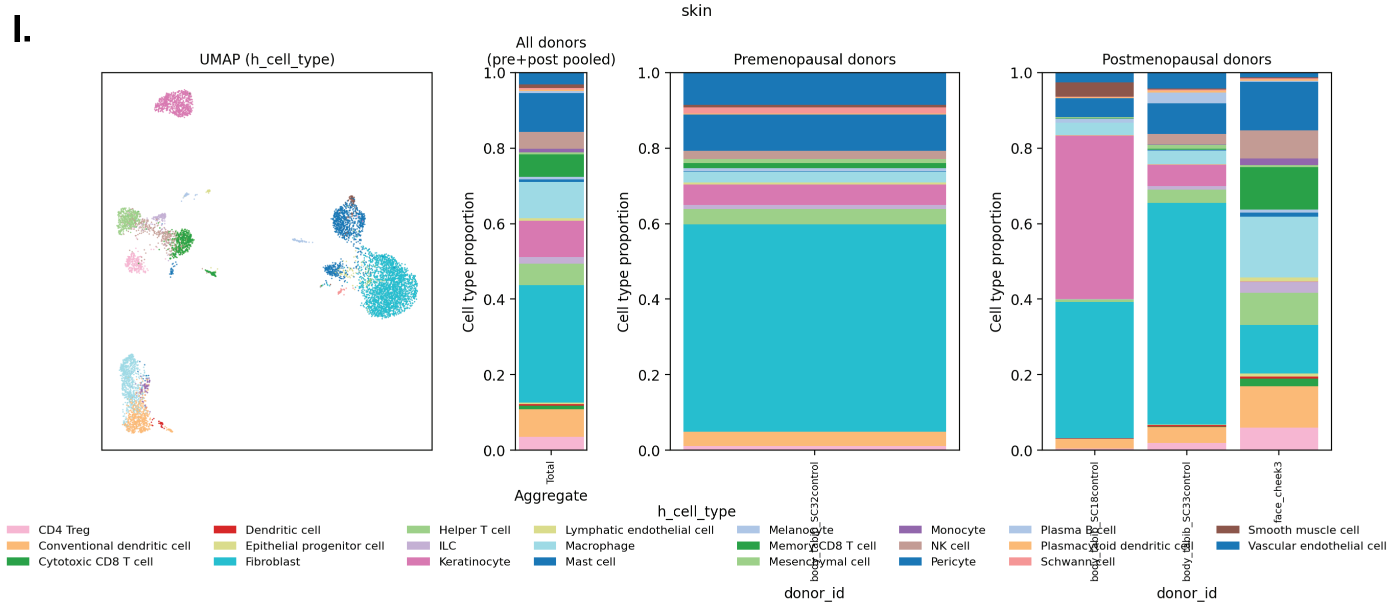
skin
UMAP (h_cell_type)
All donors
(pre+post pooled)
Premenopausal donors
Postmenopausal donors
Cell type proportion
0.0
0.2
0.4
0.6
0.8
1.0
Total
Aggregate
body_abib_SC32control
donor_id
body_abib_SC18control
body_abib_SC33control
face_cheek3
donor_id
h_cell_type
CD4 Treg
Conventional dendritic cell
Cytotoxic CD8 T cell
Dendritic cell
Epithelial progenitor cell
Fibroblast
Helper T cell
ILC
Keratinocyte
Lymphatic endothelial cell
Macrophage
Mast cell
Melanocyte
Memory CD8 T cell
Mesenchymal cell
Monocyte
NK cell
Pericyte
Plasma B cell
Plasmacytoid dendritic cell
Schwann cell
Smooth muscle cell
Vascular endothelial cell

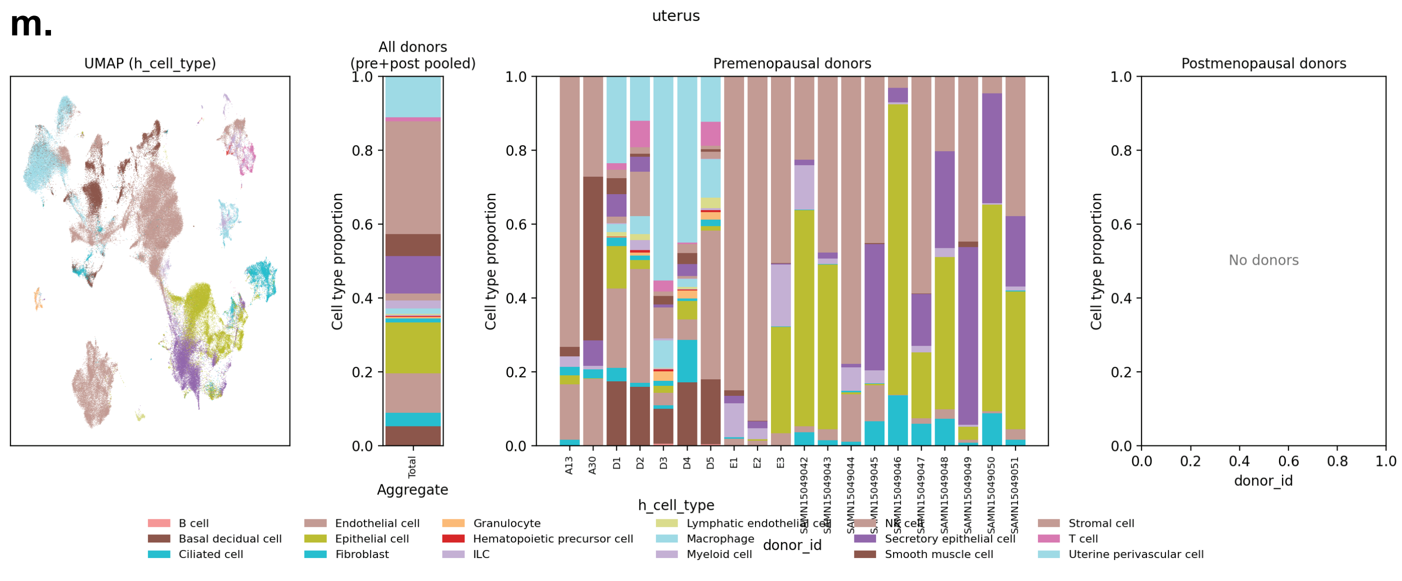


**Supplemental Figure 3. UMAPs and donor-specific cell type composition per tissue**.

(Left) UMAP; (left-middle) pooled donor cell type composition; (right-middle) individual pre-menopausal donor cell composition; (right) individual pre-menopausal donor cell composition for each organ. Organ order top-to-bottom: (a) adipose, (b) breast, (c) eye, (d) fallopian tube, (e) heart, (f) intestine, (g) liver, (h) kidney, (i) lung, (j) ovary, (k) pancreas, (l) skin, (m) uterus. Harmonized cell type labels are given in each respective sub-figure legend.

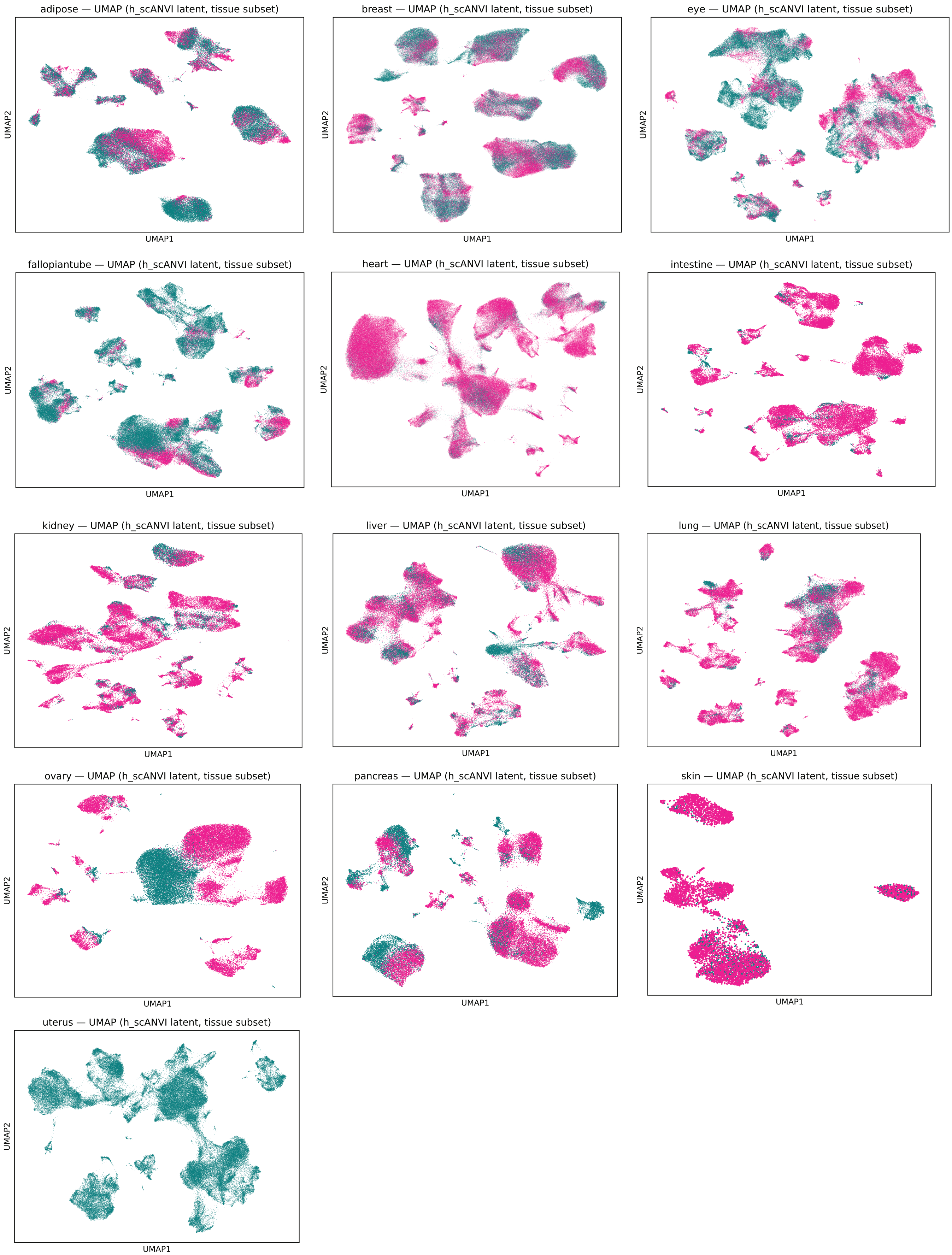


**Supplemental Figure 4. UMAPs of each organ colored by menopausal status.** UMAP projections of each organ subsetted from the full integrated MenoMap colored by menopausal status. Teal = cells from premenopausal donors; pink = cells from postmenopausal donors.

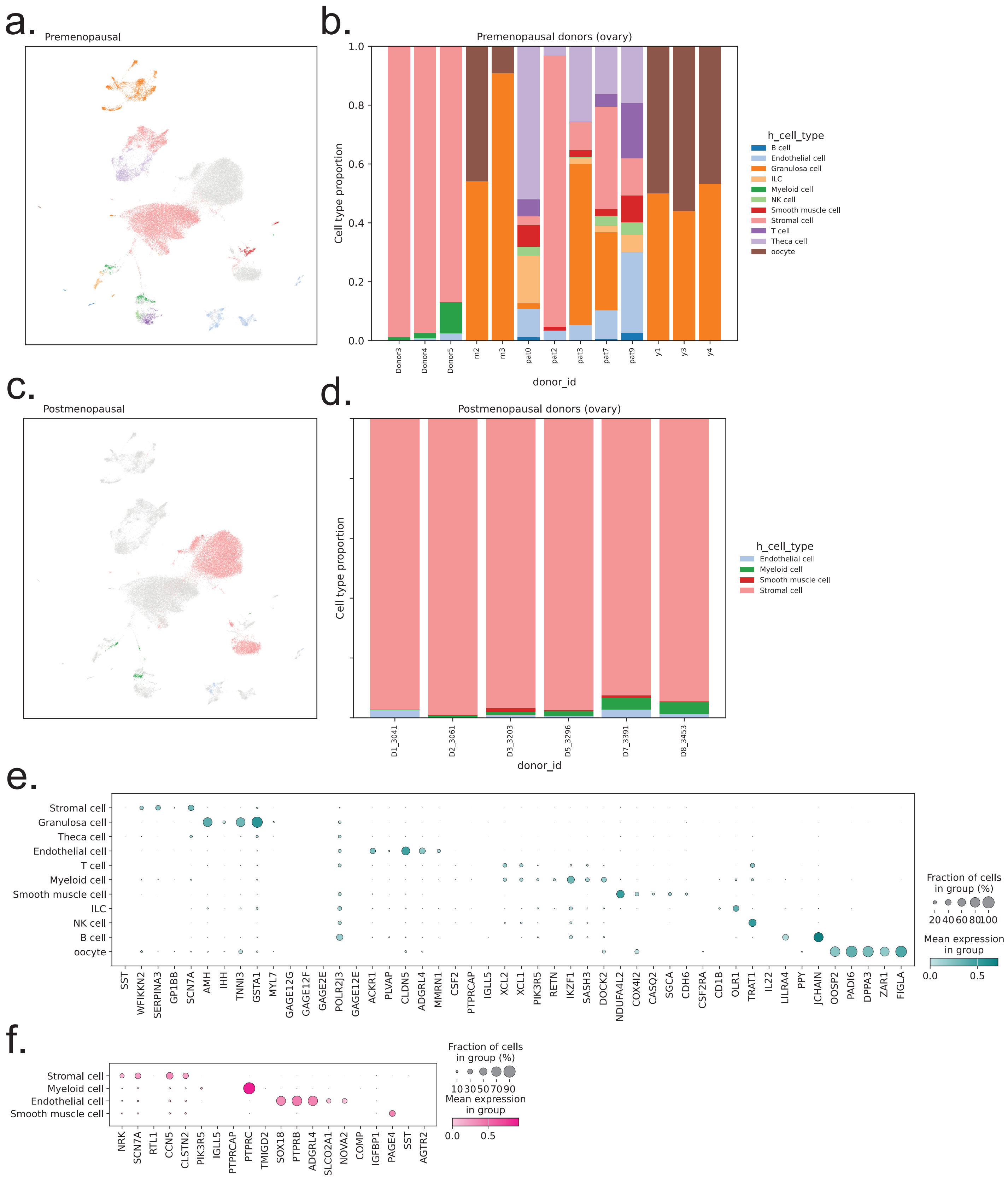


**Supplemental Figure 5. Validation of pre- and post-menopausal ovarian subsets.**

**(a)** UMAP projection of ovarian cell types colored by premenopausal cell type; postmenopausal cells colored grey.

**(b)** Ovarian cell type proportions in individual premenopausal donors.

**(c)** UMAP projection of ovarian cell types colored by postmenopausal cell type; premenopausal cells colored grey.

**(d)** Ovarian cell type proportions in individual postmenopausal donors.

**(e)** Dot plot of top ovarian cell type marker genes among premenopausal ovary donors identified by SCANVI differential expression (each ovarian cell type versus all others, mode='change'). For each cell type, the top five upregulated genes ranked by mean log fold change (lfc_mean) with FDR < 0.05 are shown. Rows correspond to tissues; dot color indicates mean log-normalized expression (normalize_total + log1p on raw counts), and dot size indicates the fraction of cells expressing each gene.

**(f)** Dot plot of top ovarian cell type marker genes among postmenopausal ovary donors identified by SCANVI differential expression (each ovarian cell type versus all others, mode='change'). For each cell type, the top five upregulated genes ranked by mean log fold change (lfc_mean) with FDR < 0.05 are shown. Rows correspond to tissues; dot color indicates mean log-normalized expression (normalize_total + log1p on raw counts), and dot size indicates the fraction of cells expressing each gene.

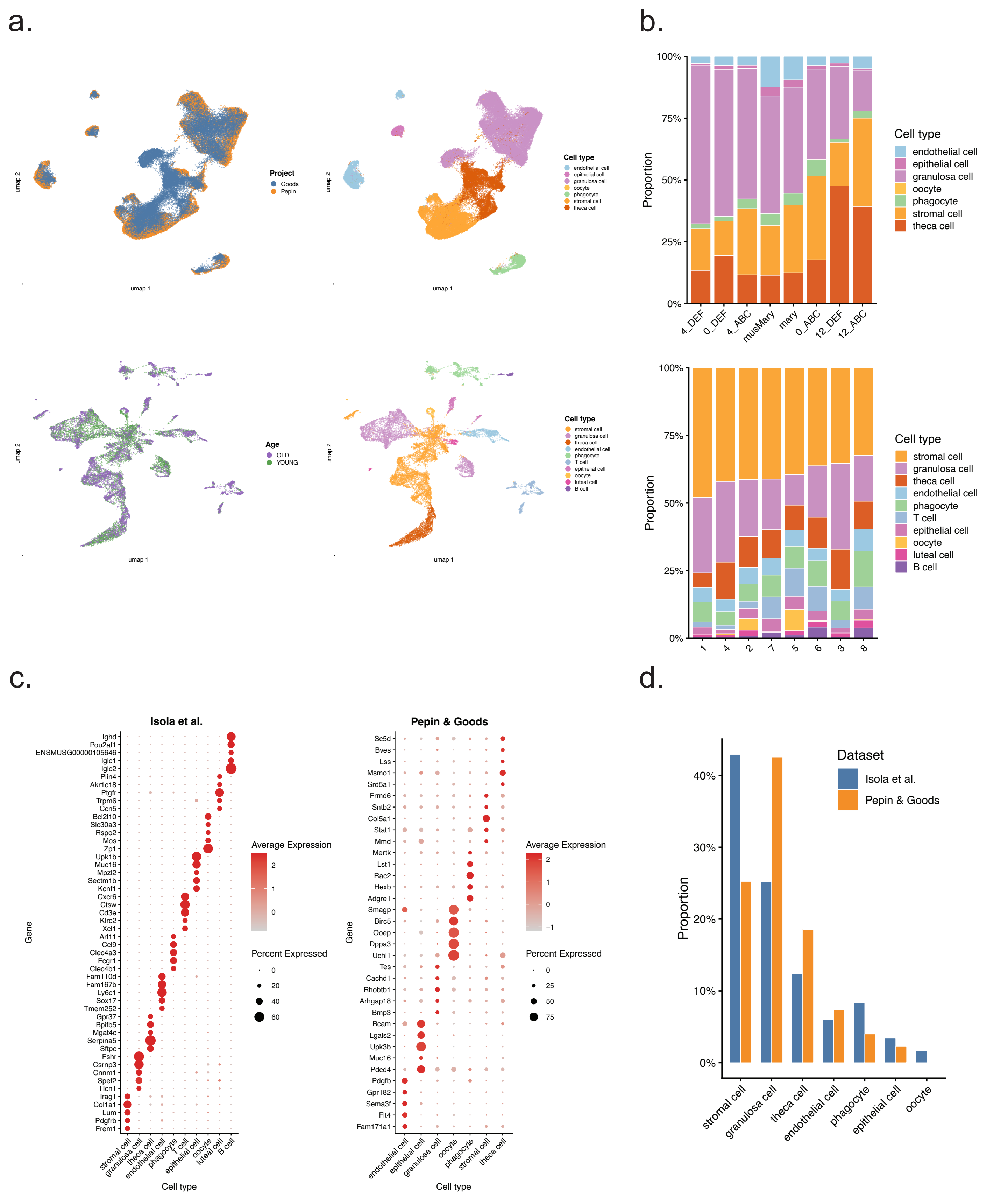


**Supplemental Figure 5. Characterization and cross-dataset validation of mouse ovarian single-cell atlases**

**(a)** UMAP projections of two independent mouse ovarian scRNA-seq datasets: the Pepin & Goods cohort colored by study of origin (left) and by annotated cell type (center-left); and the Isola et al. 2024 dataset colored by age group (center-right) and by cell type (right).

**(b)** Stacked bar charts showing the proportional cell-type composition per individual sample in the Isola et al. 2024 dataset (upper) and the Pepin & Goods dataset (lower), ordered by the most abundant cell type per sample.

**(c)** Dot plots of the top marker genes per cell type, identified by one-vs-rest Wilcoxon rank-sum tests, for the Isola et al. 2024 dataset (left) and the Pepin & Goods dataset (right). Dot size encodes the fraction of cells in the cluster expressing the gene; color intensity reflects the scaled mean expression level.

**(d)** Bar charts comparing cell-type proportions between young and old animals within the Isola et al. 2024 dataset (left) and across shared cell types between the two mouse datasets (right), illustrating cross-dataset reproducibility of cellular composition estimates.
